# Functional Spectrum of USP7 Pathogenic Variants in Hao-Fountain Syndrome: Insights into the Enzyme’s Activity, Stability, and Allosteric Modulation

**DOI:** 10.1101/2025.03.20.644318

**Authors:** Emilie J. Korchak, Mona Sharafi, Isabella Jaen Maisonet, Pilar Caro, Christian P. Schaaf, Sara J. Buhrlage, Irina Bezsonova

## Abstract

Hao-Fountain syndrome is a rare neurodevelopmental disorder caused by mutations in the de-ubiquitinating enzyme USP7 (Ubiquitin Specific Protease 7). Due to the novelty of the disease and its poorly understood molecular mechanisms, treatments for the syndrome are currently lacking. This study examines the effects of 11 patient-derived variants located within the catalytic domain of USP7, focusing on their impact on the enzyme’s activity, thermodynamic stability, and substrate recognition. Our findings reveal a spectrum of functional consequences, ranging from complete inactivation to hyperactivation of USP7. Notably, we identify a specific subset of pathogenic variants whose catalytic activity can be significantly boosted using a novel allosteric activator. These results provide the first insight into USP7 malfunction in Hao-Fountain syndrome-linked variants and pave the way for improved prognostic approaches and targeted treatments in the future.

## INTRODUCTION

Hao-Fountain syndrome (OMIM # 616863) is a rare neurodevelopmental disorder characterized by developmental delays, speech impairments, including non-verbal individuals, hypotonia, seizures, and varying degrees of intellectual disability^1,2^. Currently, there are no available treatments for this condition. The syndrome is typically caused by heterozygous *de novo* pathogenic variants or microdeletions of the *USP7* gene located on chromosome 16p13.2, which encodes the Ubiquitin-Specific Protease 7 (USP7) enzyme^1-3^. The precise molecular mechanisms underlying Hao-Fountain syndrome and the impact of these pathogenic variants on USP7’s function remain unknown.

USP7, also known as Herpesvirus-Associated Ubiquitin-Specific Protease (HAUSP), is a ∼128 kDa de-ubiquitinating enzyme (DUB) responsible for removing ubiquitin modifications from mono- and poly-ubiquitinated proteins, thereby influencing their cellular fate^4^. It is involved in several crucial cellular processes, including DNA damage response, epigenetics, immune response, signaling, and the apoptotic and p53 pathways^5-11^. Overexpression of USP7 has been associated with increased tumor aggressiveness in human prostate cancer and poor prognosis in lung squamous cell carcinoma and large cell carcinoma^12,13^, while reduced USP7 expression is linked to adenocarcinoma progression in non-small cell lung cancer^14^. Various USP7 inhibitors have been explored as potential anti-cancer treatments^5,7,15,16^ and the novel allosteric DUB activator (MS-8) specific for USP7 has been reported in the companion manuscript.

The USP7 protein is composed of seven domains: the N-terminal TRAF-like domain (Tumor necrosis factor Receptor–Associated Factor), the catalytic domain, and five ubiquitin-like (UBL) domains, followed by a regulatory C-terminal tail. The TRAF-like and UBL domains are crucial for substrate recognition and enzyme specificity^17-36^. The catalytic domain of USP7 binds the ubiquitin moiety of the ubiquitinated substrate and cleaves the isopeptide bond to remove the post-translational modification. USP7 possesses a built-in regulatory mechanism that is unique among the Ubiquitin Specific Protease family of enzymes. Substrate binding induces a conformational switch in the catalytic triad (C223, H464, D481) from an inactive to an active state^17,37^, and the association between the disordered regulatory C-terminal tail and the catalytic domain is required for the full activity^30,38-41^.

Variants associated with Hao-Fountain syndrome are dispersed throughout the USP7 protein, with a majority affecting the catalytic domain. In this study, we focus on missense variants within this domain, examining their effects on USP7’s enzymatic activity, thermodynamic stability, and ubiquitin binding. We also use the novel allosteric activator of USP7, MS-8, to assess whether the activity of disease-causing variants can be enhanced. Using this approach, we identified a group of pathogenic variants that can potentially be rescued. Our results offer the first insight into the causes of USP7 malfunction in Hao-Fountain syndrome and provide a foundation for the development of effective treatment strategies in the future.

## RESULTS

### Distribution of USP7 variants associated with Hao-Fountain syndrome

The distribution analysis of missense and nonsense mutations linked to Hao-Fountain syndrome reveals that the pathogenic variants are dispersed throughout the USP7 protein. However, the majority are found within the catalytic domain **(Figure 1A)**^3^, which harbors 19 missense mutations. Notably, several of these, including variants of Y331 and G392, have been identified in multiple individuals, suggesting their functional significance. Other hot spots of causal variants are located within the substrate-binding TRAF and UBL12 domains, suggesting that they may result in defects in the recognition of specific USP7 substrates.

**Figure 1.**
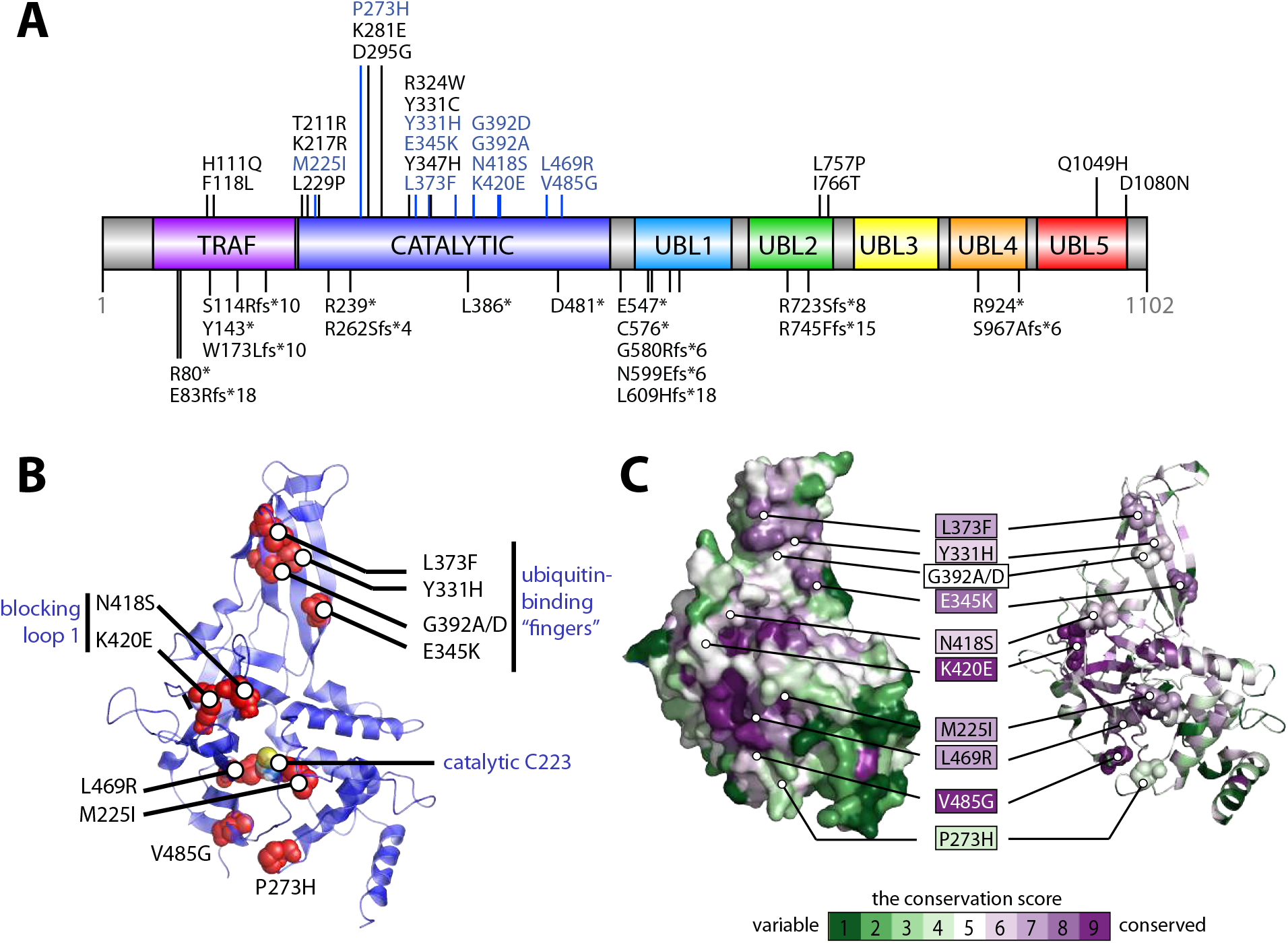
Distribution of USP7 variants associated with Hao-Fountain syndrome. **A**. The distribution of USP7 variants linked to Hao-Fountain syndrome is shown on the schematic representation of the protein drawn by DOG 2.0^51^. The USP7 structural domains, including the TRAF-like domain (TRAF), the catalytic domain (catalytic), and the ubiquitin-like domains 1 through 5 (UBL1, UBL2, UBL3, UBL4, and UBL5) are labeled. Missense and nonsense mutations are shown on the top and bottom of the protein schematic, respectively. Pathogenic variants investigated in this study are highlighted in blue and shown as red spheres mapped on the structure of the catalytic domain (PDB ID: 5JTV)^38^ in panel **B**. The catalytic cysteine of the active site (Cys223) is labeled. Clusters of mutations located on the ubiquitin-binding “fingers” and blocking loop 1 of the domain are grouped and annotated. **C**. Surface (left) and ribbon (right) representations of the USP7 catalytic domain color-coded according to its sequence conservation score calculated using the Consurf server^42-45^. The disease-linked variants are highlighted and labeled using the same color scheme, where the most variable and the most conserved residues are shown in green and purple, respectively.

For further detailed functional studies, we selected a subset of eleven missense variants located within the catalytic domain. They are highlighted in blue in **Figure 1A**. As illustrated in **Figure 1B**, these variants are spread across the catalytic domain: four are positioned near the catalytic cleft (P273H, M225I, L469R, and V485G), five are located on the ubiquitin-binding fingers (Y331H, E345K, L373F, G392A, and G392D), and two are situated on Blocking Loop 1, which gates access to the active site (N418S and K420E). Sequence conservation analysis of the catalytic domain against homologous non-redundant sequences, performed using the Consurf server^42-45^, revealed that the disease-causing variants affect evolutionarily conserved residues in USP7. As illustrated in **Figure 1C**, which maps conservation scores onto the USP7 structure, all pathogenic variants, except P273H are located in highly conserved positions, underscoring their potential functional significance.

The pathogenic variants could potentially disrupt USP7 function by compromising the structural integrity of the catalytic domain, altering its catalytic activity, or impairing its ability to bind ubiquitinated substrates. To evaluate these effects, each variant was introduced into USP7 expression plasmids encoding either the isolated catalytic domain (residues 207-560) or the full-length protein (residues 1-1102). We then assessed their impact on USP7’s thermodynamic stability, enzymatic activity, and ubiquitin-binding capabilities.

### Effect of Hao-Fountain variants on the structural integrity of USP7

To evaluate the impact of Hao-Fountain variants on the structural integrity of the isolated USP7 catalytic domain, we first established whether the domains harboring respective variants remain folded at 30°C. To test this, we isotopically labeled each domain variant and recorded their 2D ^1^H-^15^N TROSY NMR spectra. The resulting “fingerprint” spectra of 10 mutants (M225I, P273H, Y331H, E345K, L373F, G392A, G392D, N418S, K402E, V485G) displayed a pattern of well-dispersed peaks characteristic of a folded protein with minimal changes compared to the WT spectrum **(Supplementary Figure S1)**.

To further investigate the effect of the variants on protein stability, we analyzed the thermal denaturation profiles of the mutants and compared their melting temperatures (T_m_) to the wild type (WT) **(Figure 2A)**. Mutants that exhibited a decrease in the melting temperature (ΔT_m_) of more than 2°C compared to the WT were classified as destabilizing. Our analysis identified four mutants, P273H, K420E, V485G, and L469R, as structurally destabilizing. Among these, L469R had the strongest impact, suggesting severe folding defects. All mutants are color-coded in **Figure 2** and throughout according to their thermal stability. As anticipated, variants affecting surface-exposed residues, such as those on the ubiquitin-binding “fingers,” generally had minimal impact on T_m_. In contrast, variants in buried residues, such as L469R and V485G, significantly compromised the structural integrity of the enzyme **(Figure 2B-C)**.

**Figure 2.**
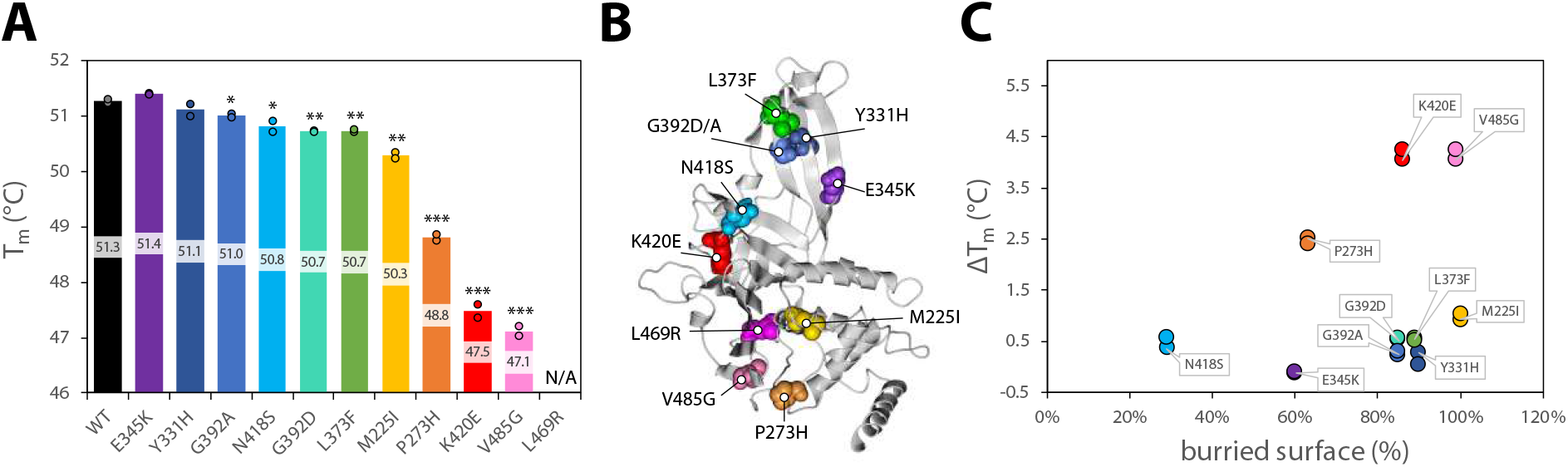
Effect of Hao-Fountain variants on the structural integrity of USP7. **A**. A bar graph displaying the melting temperatures (T_m_) of USP7 catalytic domain variants, measured using isolated catalytic domains. The bars are color-coded in a rainbow pattern, ranging from the most stable mutant (purple) to the least stable mutant (pink) marked with their individual melting temperatures. No change in intrinsic tryptophan fluorescence was observed during thermal unfolding of L469R. Dots represent individual experimental repeats. Unpaired t-tests were used to assess differences in melting temperatures between the mutants and the WT (^*^ p-value<0.05, ^**^ p-value<0.01, ^***^ p-value<0.001). **B**. Hao-Fountain variants mapped on the structure of the USP7 catalytic domain (PDB ID: 5JTV)^38^ and color-coded based on their individual thermostability. **C**. A plot of surface exposure for individual mutated residues versus the decrease in their T_m_. All data points are labeled with the name of the variant and color-coded according to their thermostability.

### Effect of pathogenic variants on the USP7 activity

To assess the impact of pathogenic variants on USP7 activity we conducted de-ubiquitination assays using purified USP7 and a minimal fluorogenic substrate, ubiquitin-AMC (7-amido-4-methylcoumarin). It has been previously reported that the FL-USP7 is significantly more active than its isolated catalytic domain due to the allosteric interaction between the domain and the regulatory C-terminus of the protein^38,39,46^. Therefore, we tested the effect of Hao-Fountain variants on both, the FL-USP7 and the catalytic domain. Consistent with previous studies, the FL-USP7 was significantly more active than the isolated catalytic domain (**Figure 3, Supplementary Figure S2** and **Table S1**).

**Figure 3.**
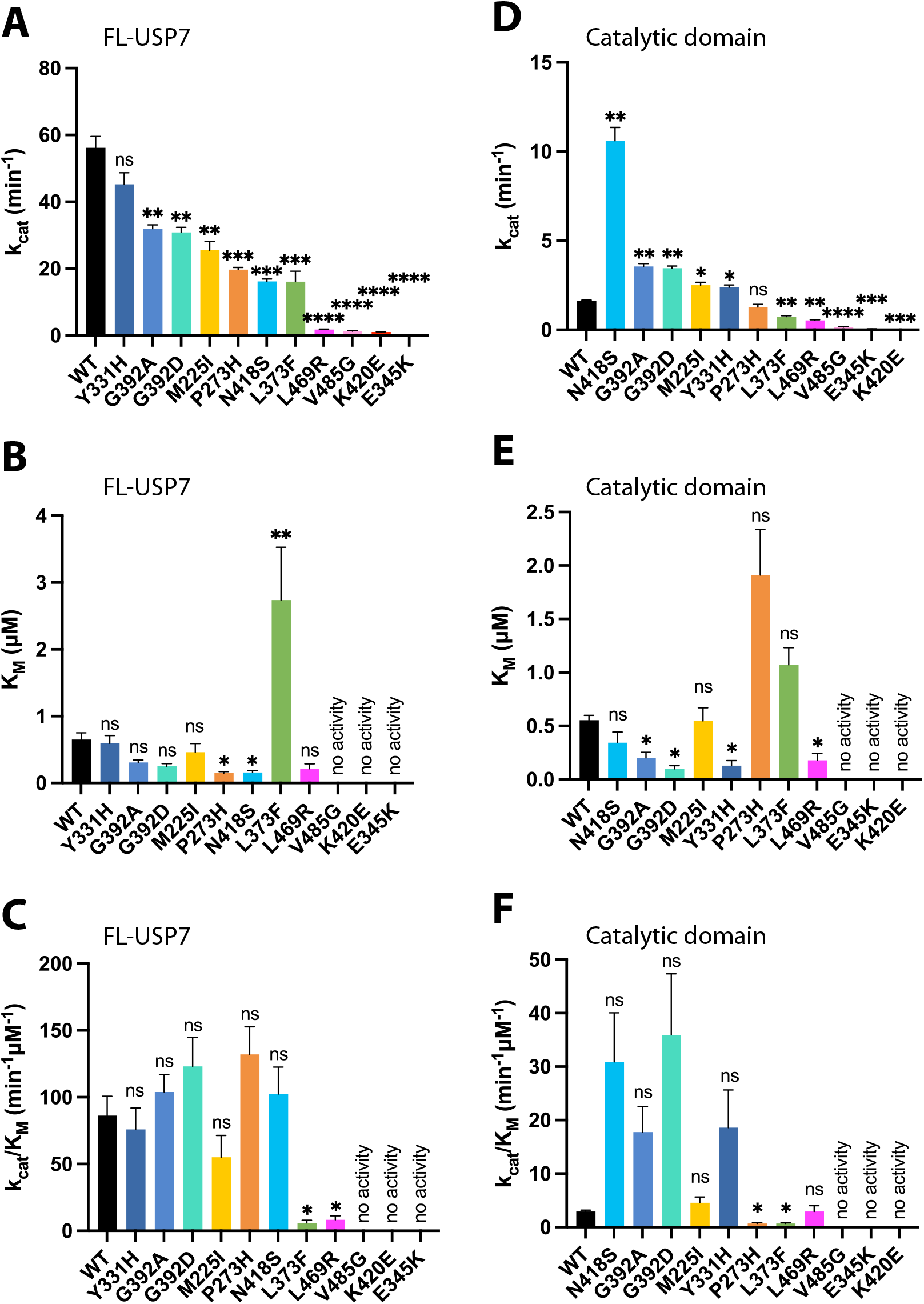
Effect of pathogenic variants on USP7 activity. Steady-state kinetic analysis of the FL-USP7 (**A-C**) and catalytic domain (**D-F**) variants measured using ubiquitin-AMC as a fluorogenic substrate. **A**. Comparison of the turnover number (k_cat_) of the FL-USP7 and its mutants. **B**. Comparison of the Michaelis-Menten constant (K_M_) of the FL-USP7 and its mutants. **C**. Comparison of the catalytic efficiency (k_cat_/K_M_) of the FL-USP7 and its mutants. **D**. Comparison of the turnover number (k_cat_) of the isolated catalytic domain and its mutants. **E**. Comparison of the Michaelis-Menten constant (K_M_) of the isolated catalytic domain and its mutants. **F**. Comparison of the catalytic efficiency (k_cat_/K_M_) of the isolated catalytic domain and its mutants. The number of repeats was n=2 for all FL proteins, except WT n=7 and V485G n=4. The number of repeats was n=2 for all catalytic domain proteins, except N418S, M225I and V485G, where n=4. Unpaired t-tests were used to assess differences in kinetic parameters between the mutants and the WT (ns=non significant, ^*^ p-value<0.05, ^**^ p-value<0.01, ^***^ p-value<0.001, ^****^ p-value<0.0001). All bars are color-coded according to the thermostability of each variant. Also see Supplementary **Figure S2**.

The enzyme kinetics analysis revealed that in the context of the FL-USP7, nearly all mutations resulted in loss of activity (**Figures 3A, C**) as evidenced by their lower turnover numbers (k_cat_) compared to the WT. The impact of individual mutations ranged from a 20% reduction in k_cat_ to nearly complete inactivation of the enzyme (**Figures 3A, Supplementary Table S1**). As expected, the structurally unstable mutants, such as K420E, L469R, and V485G, exhibited nearly complete loss of activity characterized by very low k_cat_, whereas the k_cat_ of the stable mutants, such as Y331H, were comparable to the WT. Interestingly, despite their structural stability, E345K and L373F mutants experienced a dramatic loss in activity. The significant increase in Michaelis-Menten constant (K_M_) for L373F (**Figure 3B**) suggests that its decreased catalytic efficiency is primarily caused by defects in substrate binding. Similar to L373F, the E345K variant, located on the ubiquitin-binding fingers (**Figure 1B**), likely causes a similar defect; however, its K_M_ value could not be estimated due to its complete lack of activity.

In contrast to the FL-USP7, the effects of the disease-causing variants on the isolated catalytic domain varied from complete inactivation to unexpected hyperactivation (**Figures 3D, F, Supplementary Table S1**). Consistent with the full-length enzyme data, variants that caused folding defects (L469R, V485G, and K420E) also resulted in an inactive isolated catalytic domain. However, variants such as N418S, G392A, and G392D led to significant enhancement of the enzyme activity, with up to a 7-fold increase in k_cat_. The N418S pathogenic variant, in particular, caused the most substantial enhancement. Located on a dynamic loop that regulates access to the catalytic cleft, this variant likely enhances the enzyme’s activity by improving accessibility to the active site. Interestingly, a decrease in K_M_ values measured for the hyperactive G392A, G392D, and Y331H variants suggests that their enhanced activity is caused by an improved affinity for the substrate compared to the WT (**Figure 3E**), which is consistent with their location next to each other on the adjacent β-strands of the ubiquitin-binding fingers (**Figure 1B**). Increased K_M_ values have been measured for pathogenic variants L373F and P273H (**Figure 3E**), pointing to defects in substrate recognition, which lead to their reduced catalytic efficiency (k_cat_/K_M_). Comparative analysis of the k_cat_/K_M_ values of all variants revealed that structurally unstable variants and variants causing substrate-recognition defects generally result in enzyme inactivation. In contrast, the effect of remaining pathogenic variants varies from WT-like activity to hyperactivation of the isolated catalytic domain.

Overall, a consistent pattern of the functional impact of the disease-causing mutations on the activity of the full-length enzyme and its isolated catalytic domain has been observed. However, the hyperactive mutations detected in the isolated catalytic domain did not have such a dramatic effect in the FL-USP7. This suggests that despite being active in the context of the isolated domain, some variants may still interfere with the full enzyme activity, which requires an allosteric intramolecular interaction between the catalytic domain and the regulatory C-terminus. These results further highlight the complex regulatory mechanism of USP7 activity.

### Effect of pathogenic variants on USP7 affinity to ubiquitin

USP7 interacts with and de-ubiquitinates a multitude of diverse targets. While specific substrates are recognized by the TRAF and UBL domains of the enzyme, their ubiquitin modifications are recognized by the catalytic domain. To test the impact of the Hao-Fountain variants on USP7’s affinity to ubiquitin, we used NMR chemical shift perturbation (CSP) assays, which are best suited for quantifying weak interactions typical for ubiquitin. In these assays, ^15^N-labeled catalytic domains carrying a causal variant were titrated with unlabeled ubiquitin and their dissociation constants (K_D_) were determined from the resulting CSPs **(Supplementary Figure S3)**. A comparison of the individual K_D_ values identified E345K as a mutant, which is severely deficient in ubiquitin-binding **(Figure 4A)**.

**Figure 4.**
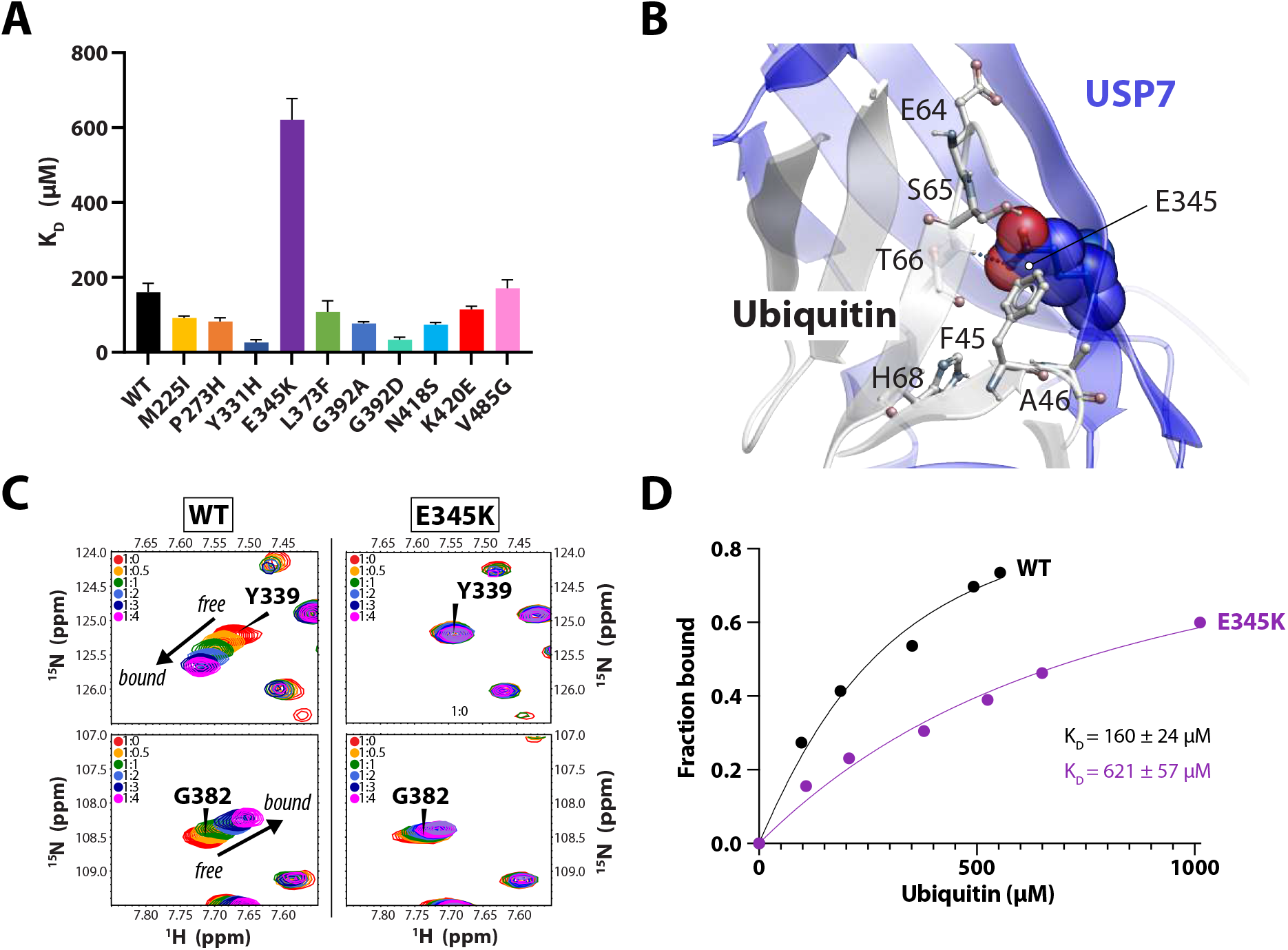
Effect of pathogenic variants on USP7 affinity to ubiquitin. **A**. Comparison of ubiquitin-binding affinities of the USP7 catalytic domain and its mutants. The individual dissociation constants (K_D_) were measured using NMR chemical shift perturbations in the ^15^N TROSY spectra of the ^15^N-labeled domain upon the addition of unlabeled ubiquitin. All data are color-coded according to the thermostability of each variant. **B**. The location of E345 on the USP7/ubiquitin binding interface (PDB ID: 1NBF)^17^. USP7 is shown in blue, ubiquitin in grey, and both molecules are labeled. **C**. Select regions of ^15^N TROSY spectra of the WT (left panels) and the ubiquitin-binding deficient mutant E345K (right panels) show chemical shift perturbations (Δω) of residues Y339 and G382. These residues are located on the ubiquitin-binding “fingers” of the catalytic domain and demonstrate changes from free to ubiquitin-bound states of the protein. **D**. Plot showing the normalized global Δω measured for the WT and E345K catalytic domains as a function of ubiquitin concentration, used to estimate the binding affinities. Also see Supplementary **Figure S3** for all other mutants.

The E345K mutant is structurally stable but catalytically inactive as seen in **Figure 2A** and **Figure 3**. The variant is located on the “ubiquitin-binding fingers” of the enzyme **(Figure 4B)**. Therefore, it is expected that the charge-reversing mutation E345K may disrupt the interaction with ubiquitin. Indeed, in contrast to the WT, the NMR binding assays show minimal peak perturbations in the spectrum of E345K mutant upon titration with ubiquitin **(Figures 4C and Supplementary Figure S3E)**, resulting in an estimated K_D_ of ∼600 μM **(Figure 4D)**. Taken together, this data suggests that the complete loss of the E345K mutant’s activity is caused by its inability to effectively interact with ubiquitin.

Surprisingly, another inactive “fingers” mutant, L373F, bound ubiquitin with the K_D_ similar to the WT. However, significantly fewer peaks were perturbed in its NMR spectrum during the titration, suggesting a reduced binding interface. These results are in good agreement with the increased Michaelis-Menten constant, which suggests a sub-optimal substrate affinity of L373F compared to the WT **(Figures 3B and E)**.

In contrast, variants Y331H and G392A/D also found in the “fingers” region, increased USP7’s affinity for ubiquitin **(Figure 4A)**. This enhancement likely contributes to the hyperactivation of the catalytic domain observed in **Figures 3D-F**.

### Activation of Hao-Fountain syndrome variants by an allosteric activator

In the companion manuscript, we have shown that WT USP7 can be activated by an allosteric compound MS-8. This small molecule effectively mimics the regulatory C-terminal tail of the enzyme by targeting the allosteric site on the catalytic domain. To assess whether the activator can enhance the activity of Hao-Fountain-associated variants, we treated the full-length enzymes with increasing concentrations of MS-8 and measured their activity using DUB assays **(Figure 5 and Supplementary Figure S4)**. The enzyme kinetics analyses revealed that the initial reaction rate (V_0_) of most mutants can be enhanced by MS-8 to the WT level and beyond in a concentration-dependent manner **(Figure 5A)**. As anticipated, the activity of structurally unstable mutants like L469R and K420E, as well as mutants with impaired ubiquitin-binding, such as E345K and L373F, could not be rescued **(Figure 5B)**. This indicates that allosteric activation is insufficient to compensate for defects in protein folding or substrate recognition. On the other hand, mutants retaining the catalytic activity could be further activated.

**Figure 5.**
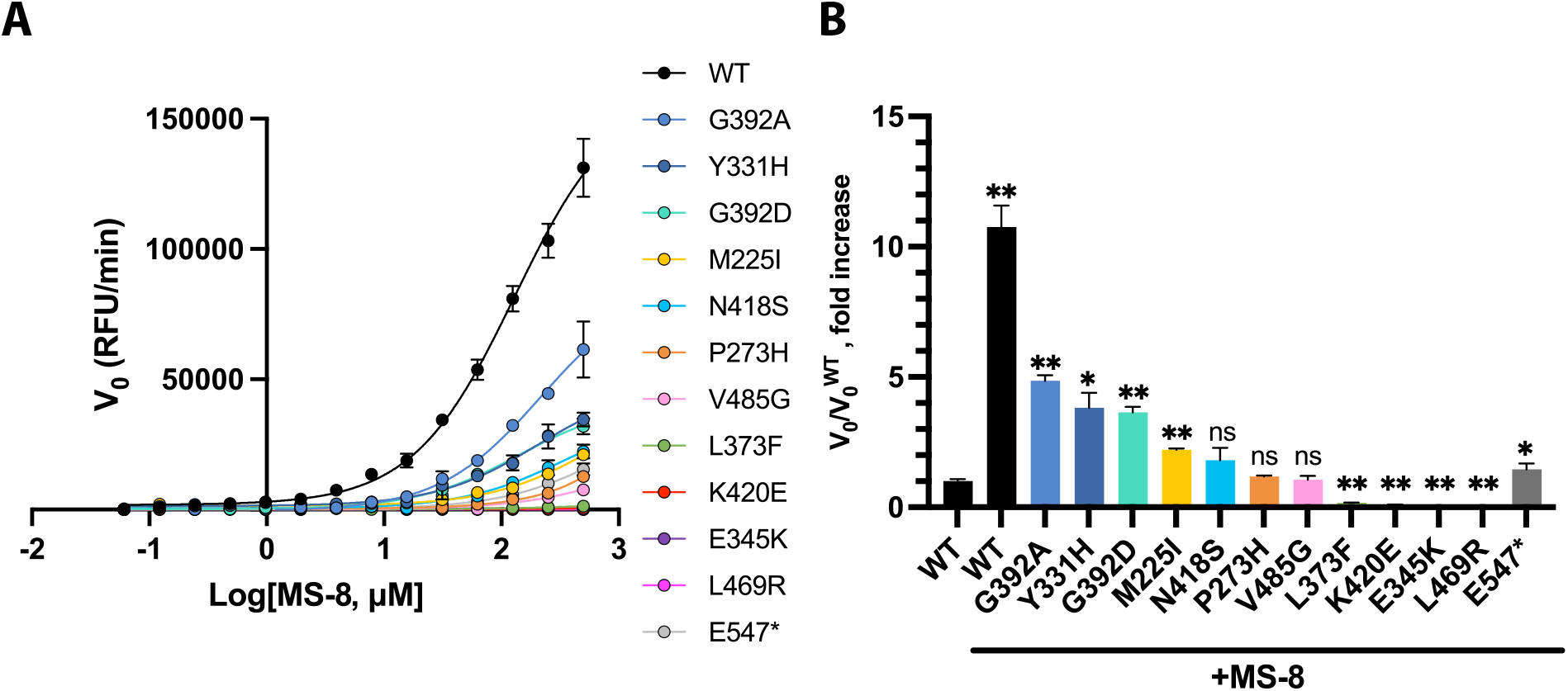
Enhancement of pathogenic variants’ activity by MS-8. **A**. Concentration-dependent enhancement of deubiquitinating activity in FL-USP7 and its mutants by MS-8 using ubiquitin-rhodamine as the fluorogenic substrate. The initial velocity (V_0_) is plotted against log[MS-8]. **B**. Initial velocity enhancement caused by the treatment of the mutants with 250 μM MS-8 relative to the velocity of the untreated WT (V_0/V0_^WT^). The number of repeats was n=2 for all proteins except E547*, where n=6. Unpaired t-tests were used to assess differences in V_0_ fold increase between the mutants and the WT (ns=non significant, ^*^ p-value<0.05, ^**^ p-value<0.01). All data are color-coded according to the thermostability of each variant.

Since the activator interacts with the enzyme’s allosteric site, which is normally occupied by its C-terminus, we reasoned that the pathogenic variants resulting in C-terminally truncated USP7 with the intact catalytic domain **(Figure 1A)** might be effective targets for re-activation with MS-8. In Hao-Fountain syndrome patients, such variants include E547^*^, C576^*^, G580Rfs^*^6, N599Efs^*^6, L609Hfs^*^18, R723Sfs^*^8, R745Ffs^*^15, R924^*^ and S967Afs^*^6. Among these, E547^*^ causes the largest truncation. Therefore, we chose it as the representative pathogenic variant to test our hypothesis. Indeed, its treatment with 250 μM MS-8 resulted in a 47-fold increase in initial velocity, effectively restoring its activity to the WT level **(Supplementary Figure S4** and **Figure 5B)**. This suggests that the activity of other C-terminally truncated mutants can also be rescued.

These findings strongly suggest that allosteric activation is a promising strategy for enhancing the activity of USP7 and some of its disease-causing mutants.

## DISCUSSION

USP7, a vital deubiquitinating enzyme, is integral to numerous cellular processes by removing ubiquitin tags from substrates, thus influencing their stability and function^4^. Its role extends across critical biological pathways, including DNA damage repair, immune response modulation, and various signaling mechanisms^5-11^. Given the extensive functions of USP7, precise regulation of its activity is essential for maintaining cellular health. Disruptions, such as overexpression of USP7, have been linked to poor cancer prognosis, highlighting the significance of its regulation in cellular integrity^12,13^.

The involvement of USP7 in the endosomal protein recycling pathway led to the identification of Hao-Fountain syndrome in 2015 with only six patients. The number of cases has been steadily increasing due to advancements in genetic testing and growing awareness, reaching a total of 250 cases worldwide to date^47^. Within this pathway, USP7 acts as a molecular rheostat in conjunction with MAGE-L2 and the E3 ligase TRIM27, facilitating WASH-dependent endosomal protein recycling^1^. Notably, mutations and deletions in MAGE-L2 have been implicated in Schaaf-Yang and Prader-Willi syndromes, both of which are associated with neurodevelopmental challenges^48-50^. USP7 mutations result in similar phenotypes that include intellectual disability, autism spectrum disorder, epilepsy, and hypogonadism. Considering the significant overlap in symptoms and the limited understanding of USP7 dysregulation, we aimed to investigate the functional effects of USP7 pathogenic variants *in vitro*. A comprehensive understanding of how these variants impact USP7 is crucial for identifying potential therapeutic approaches in the future.

Our study examined the functional consequences of 11 missense variants within the catalytic domain of USP7 and tested whether their activity can be rescued using a novel allosteric activator of USP7 as a tool. We have shown that the mutations impact USP7’s function in three primary ways: altering the structural integrity of the protein, disrupting ubiquitinated substrate recognition, and, ultimately, affecting its enzymatic activity. Remarkably, the allosteric activator, MS-8, could effectively enhance the catalytic activity of all mutants except for those with severe structural defects that impact either the enzyme’s folding or its ability to bind ubiquitin. Furthermore, in agreement with its allosteric mode of action, MS-8 effectively restored the activity of truncating variants lacking the regulatory C-terminal tail.

Although some pathogenic variants irreversibly lost their activity, we have shown that MS-8 could also enhance the activity of the WT enzyme. Since Hao-Fountain syndrome is caused by heterozygous mutations in the *USP7* gene, conceivably, activation of the WT USP7 could compensate for the non-activatable mutant copy.

A correlation between specific disease-causing variants and the disease phenotype is key for understanding the mechanism of pathogenesis and the development of prognostic and treatment strategies in the future. A recent study introduced a severity score specific to Hao-Fountain syndrome and showed that pathogenic variants located in the catalytic domain of the USP7 protein are associated with more severe outcomes^3^. Here, we provided a detailed functional characterization of the catalytic domain variants. Ultimately, our data revealed two groups of mutations: a group that irreversibly lost its catalytic activity due to defects in protein folding and substrate binding and a group with altered activity.

The K420E variant resulted in a complete loss of function in both the catalytic domain and the FL-USP7. In agreement with this, the patient carrying the K420E mutation has a high severity score^3^ and suffers from severe general developmental delays leading to motor and speech dysfunction, intellectual disability, hypotonia, eye anomalies, obesity, and an abnormal threshold for pain.

In contrast, G392A and G392D variants caused an overall enzyme efficiency increase **(Figure 3C** and **F)**, although their impact on the isolated catalytic domain was more pronounced compared to the FL-USP7. We expected that a more active USP7 variant would result in a less severe phenotypical outcome. Surprisingly, both patient’s phenotypes are more severe than what has been reported for individuals with other USP7 mutations. A high severity score of 0.76 and 0.87 was assigned to G392A and G392D respectively^3^. Both variants share similar phenotypes including autism spectrum disorder, severe motor developmental and speech delays, hypotonia, severe motor dysfunction, speech dysfunction, eating complications, eye anomalies, and an abnormal threshold for pain. Distinctively associated with a higher severity score, G392D also manifests ADHD, contractures, chronic gastrointestinal issues, endocrine disruption, abnormal brain MRI findings, obesity, and variations in size for gestational age. In contrast, G392A is characterized by congenital anomalies. This is an intriguing distinction, as the same residue is impacted differently by each mutation.

Taken together, these findings suggest that both loss-of-function and gain-of-function mutations can result in the severe phenotype of Hao-Fountain syndrome. The investigation of larger cohorts will be necessary to better understand genotype-phenotype correlations. It is conceivable that using specific inhibitors and activators of USP7 could offer promising personalized treatment strategies for Hao-Fountain syndrome in the future. This study offers the first insights into the functional implications of the pathogenic USP7 variants and establishes a foundation for their characterization in cellular contexts. Additionally, it sets the stage for developing cell and animal models necessary to evaluate the efficacy of small molecule modulators in restoring normal USP7 function.

In summary, our comprehensive study of the functional impact of pathogenic variants associated with Hao-Fountain syndrome underscores the complex interplay between structural integrity, enzymatic activity, and allosteric modulation of the USP7 enzyme. Understanding these relationships is pivotal in advancing our understanding of USP7’s role in neurodevelopmental disorders, laying the groundwork for future development of targeted treatments and improving patient outcomes.

## RESOURCE AVAILABILITY

All plasmids for bacterial expression of the WT and mutant USP7 genes were deposited to Addgene database including the isolated catalytic domain plasmids 227201 (WT), 227202 (N418S), 227203 (G392D), 227204 (G392A), 227205 (Y331H), 227206 (M225I), 227207 (L373F), 227208 (P273H), 227209 (V485G), 227210 (K420E), 227211 (E345K), 227212 (L469R), and the FL-plasmids 227214 (WT), 227215 (N418S), 227216 (G392D), 227217 (G392A), 227218 (Y331H), 227219 (M225I), 227220 (L373F), 227221(P273H), 227222 (V485G), 227223 (K420E), 227224 (E345K), 227225 (L469R), 229001 (E547*).

## ABBREVIATIONS

USP: ubiquitin-specific protease.
DUB: deubiquitinating enzyme
TRAF: tumor necrosis factor receptor-associated factor
UBL: ubiquitin-like
WT: wild type
FL: full-length
Ub: ubiquitin
Ub-AMC: ubiquitin-7-amido-4-methylcoumarin
Ub-Rho: Ubiquitin-Rhodamine

## ACKNOWLEDGEMENTS

We would like to thank Dr. Milka Kostic for the valuable discussions of the manuscript.

## AUTHOR CONTRIBUTIONS

E.J.K. and I.B. wrote the manuscript and prepared figures. E.J.K. designed and performed all assays. E.J.K. designed and generated all USP7 mutants and purified all proteins. I.J.M., M.S. and S.J.B. provided MS-8. P.C. and C.P.S. provided lists of Hao-Fountain syndrome mutations and description of patient phenotypes. I.B. and S.J.B. conceptualized the project, acquired funding, reviewed and edited the manuscript. All authors read and approved the final version prior to submission.

## SUPPLEMENTARY FIGURE LEGENDS

**Table S1. Summary of enzyme kinetics and ubiquitin-binding affinities of USP7 mutations associated with Hao-Fountain syndrome**.

**Figure S1. NMR spectra of USP7 variants associated with Hao-Fountain syndrome**.

2D ^15^N TROSY spectra of ^15^N-labeled USP7 catalytic domain and its Hao-Fountain syndrome variants. Individual spectra are labeled with the corresponding variant name.

**Figure S2. Effect of Hao-Fountain syndrome variants on USP7 activity**.

Michaelis-Menten plots of the initial velocity (V_0_) as a function of ubiquitin-AMC concentration shown for (**A**) USP7 catalytic domain and its mutants and (**B)** FL-USP7 and its mutants.

**Figure S3. Ubiquitin binding to catalytic domains of USP7**.

**Top:** ^15^N TROSY spectra of the ^15^N-labeled USP7 catalytic domain and its mutants gradually titrated with unlabeled ubiquitin. Residue A381 is showcased for each spectrum. USP7:ubiquitin molar ratios are shown.

**Bottom:** Plot showing the global chemical shift perturbations (Δω) in the spectra as a function of ubiquitin concentration, used to estimate the binding affinities for each USP7 variant (K_D_).

**Figure S4. MS-8 enhances the activity of USP7 variants**.

Comparison of the time course of deubiquitination reaction for FL-USP7 variants alone (red) and treated with 250 μM MS-8 (blue). The untreated WT curve is shown for reference (black). 0.1 nM USP7 was used with 500 nM ubiquitin-rhodamine as its fluorogenic substrate.

## STAR METHODS

| REAGENT/RESOURCE | SOURCE | IDENTIFIER |
| --- | --- | --- |
| **Bacterial and virus strains** |  |  |
| Escherichia coli BL21(DE3) | EMD Millipore | Cat# 70235 |
| **Chemicals, peptides, and recombinant proteins** |  |  |
| Ubiquitin-7-amino-4-methylcoumarin (Ub-AMC) | SouthBay Bio | Cat# SBB-PS0043 |
| Ubiquitin-Rhodamine 110 (Ub-Rho) | UBP Bio | Cat# M3020, 3021, 3022 |
| MS-8 | This paper | N/A |
| **Recombinant DNA** |  |  |
| WT USP7 catalytic domain | This paper | Addgene 227201 |
| N418S USP7 catalytic domain | This paper | Addgene 227202 |
| G392D USP7 catalytic domain | This paper | Addgene 227203 |
| G392A USP7 catalytic domain | This paper | Addgene 227204 |
| Y331H USP7 catalytic domain | This paper | Addgene 227205 |
| M225I USP7 catalytic domain | This paper | Addgene 227206 |
| L373F USP7 catalytic domain | This paper | Addgene 227207 |
| P273H USP7 catalytic domain | This paper | Addgene 227208 |
| V485G USP7 catalytic domain | This paper | Addgene 227209 |
| K420E USP7 catalytic domain | This paper | Addgene 227210 |
| E345K USP7 catalytic domain | This paper | Addgene 227211 |
| L469R USP7 catalytic domain | This paper | Addgene 227212 |
| WT USP7 FL | This paper | Addgene 227214 |
| N418S USP7 FL | This paper | Addgene 227215 |
| G392D USP7 FL | This paper | Addgene 227216 |
| G392A USP7 FL | This paper | Addgene 227217 |
| Y331H USP7 FL | This paper | Addgene 227218 |
| M225I USP7 FL | This paper | Addgene 227219 |
| L373F USP7 FL | This paper | Addgene 227220 |
| P273H USP7 FL | This paper | Addgene 227221 |
| V485G USP7 FL | This paper | Addgene 227222 |
| K420E USP7 FL | This paper | Addgene 227223 |
| E345K USP7 FL | This paper | Addgene 227224 |
| L469R USP7 FL | This paper | Addgene 227225 |
| E547* USP7 FL | This paper | Addgene 229001 |
| **Software and algorithms** |  |  |
| Prism 10.3 | GraphPad Software Inc | https://www.graphpad.com/scientific-software/prism/ |
| PyMOL | Schrodinger | https://pymol.org/ |
| NMR Pipe | Delaglio et al., 1995 | https://spin.niddk.nih.gov/bax/software/NMRPipe/ |
| DOG 2.0 | Ren et al., 2009 | DOG 2.0 - Protein Domain Structure Visualization |
| ICM-Pro | Molsoft | http://www.molsoft.com/icm_pro.html |
| NMRfam-Sparky | Lee et al., 2014 | https://www.cgl.ucsf.edu/home/sparky/ |
| ConSurf | Yariv et al., 2023<br>Glaser et al., 2003<br>Ashkenazy et al., 2010<br>Ashkenazy et al., 2016 | http://consurf.tau.ac.il |

### Protein expression and purification

Plasmids for bacterial expression of USP7 and its mutants contained either the coding sequence of the catalytic domain of USP7 (207-560) or the FL-USP7 (1-1102) inserted into pET28a-LIC vector (Structural Genomics Consortium), downstream of the N-terminal 6xHis tag and thrombin cleavage site. Hao-Fountain syndrome mutations were introduced using site-directed mutagenesis. The constructs were sequence-verified and deposited to Addgene database.

USP7 constructs were transformed into Escherichia Coli BL21 (DE3) cells and expressed in 1L Luria Broth (LB) media for enzymatic and thermostability assays. For NMR titration experiments, ^15^N-labeled proteins were expressed in M9 minimal media with ^15^NH_4_Cl as the sole nitrogen source. Cells were grown at 37°C until OD6_00_ reached 0.8-1.0 o.u. Protein expression was induced with 1 mM IPTG at 20°C overnight. Cells were harvested by centrifugation, resuspended in lysis buffer (20 mM sodium phosphate pH 8.0, 250 mM NaCl, 5 mM imidazole, 0.5 mM PMSF), and lysed by sonication. The lysate was centrifuged at 15,000 rpm for 45 minutes, and the supernatant was filtered and applied to TALON HisPur cobalt resin (Thermo Scientific). USP7 constructs were eluted with a buffer containing 20 mM sodium phosphate pH 8.0, 250 mM NaCl, and 250 mM imidazole. Thrombin was added to remove the 6-His tag, and the samples were buffer-exchanged overnight at 4°C into 20 mM sodium phosphate pH 7.4, and 250 mM NaCl to remove imidazole and allow for cleavage. Proteins were then purified by size-exclusion chromatography using a HiLoad Superdex 200 or 75 columns (GE Healthcare) in a buffer containing 20 mM Tris pH 8.0, 250 mM NaCl, and 2 mM dithiothreitol (DTT).

### Thermostability assays

For thermal denaturation studies of the WT USP7 catalytic domain and each USP7 catalytic domain mutant, 200 µL samples at 1 µM concentration were prepared in a buffer containing 20 mM Tris pH 8.0, 250 mM NaCl, and 2 mM dithiothreitol (DTT). Each sample was loaded into UV capillaries, and thermal denaturation was conducted using the NanoTemper Tycho NT.6 (NanoTemper). The temperature was ramped from 35°C to 95°C at approximately 20°C/min. Protein unfolding was monitored using intrinsic tryptophan fluorescence, measured at 330 and 350 nm. The unfolding profiles were displayed as the brightness ratio of 350 nm/330 nm, and the first derivative of these fluorescence ratios was used to determine the inflection temperature (T_i_) or melting temperature (T_m_), which corresponds to the peak of the transition from folded to unfolded states.

### De-ubiquitination assays

Enzyme activity for WT and mutant USP7 proteins (both catalytic domain and full-length) was measured using 7-amido-4-methylcoumarin (AMC) fluorescence following cleavage of the quenched substrate ubiquitin-AMC (Ub-AMC). Reactions were conducted at 25°C in a buffer containing 20 mM Tris pH 8.0, 0.05% CHAPS, and 10 mM BME, in Corning black 384-well plates with a volume of 25 µL per well.

For full-length USP7, data were collected at 3-minute intervals over 50 minutes, and for the catalytic domain, at 10-minute intervals over 2.5 hours, using a Molecular Devices SpectraMax M3 Fluorescence Microplate Reader with excitation and emission wavelengths of 360 nm and 465 nm, respectively. Reactions were performed with constant USP7 concentrations (1 nM) and varying Ub-AMC concentrations from 0 to 2 µM for full-length and 0 to 3 µM for the catalytic domain.

Raw fluorescence data were converted to product concentration using a standard curve of AMC. The slopes of the product concentrations were analyzed in GraphPad Prism (version 10.3.1 for Mac OS X, GraphPad Software, Boston, Massachusetts USA, www.graphpad.com), where the Michaelis-Menten equation was used to model the data with non-linear regression, to plot initial velocity against Ub-AMC concentrations, and to determine steady-state kinetic parameters. Each experiment was repeated at least twice.

### NMR studies

NMR data were collected on an 800 MHz Agilent VNMRS spectrometer equipped with a cryoprobe at 30°C. Data processing was performed with NMRPipe^52^, and analysis was done using Sparky^53^. The USP7 catalytic domains and ubiquitin were in a buffer containing 20 mM Tris pH 7.5, 100 mM NaCl, 2 mM DTT, and 10% D2O (v/v).

To study ubiquitin binding, unlabeled ubiquitin was gradually added to ∼250 µM ^15^N-labeled USP7 catalytic domain up to 1:4 molar excess of ubiquitin. Chemical shift changes were monitored using 2D ^1^H-^15^N TROSY spectra across six titration points. The dissociation constant (K_D_) for the USP7/ubiquitin complex was determined by nonlinear least-squares fitting of total chemical shift changes (Δ*ω*_*obs*_) versus ligand concentration, using the following equation:

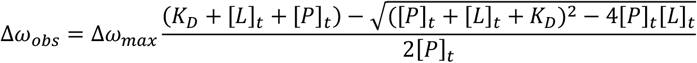

where [*P*]_*t*_ and [L]_*t*_ are the total protein and ligand concentrations, Δ*ω*_*max*_ is the chemical shift difference at saturation and Δ*ω*_*obs*_ is the total observed chemical shift change calculated as a sum of chemical shift changes of individual peaks. The fitting was performed using GraphPad Prism.

### Activity Enhancement Assays

Activity enhancement assays for the WT and mutant USP7 were performed using full-length enzymes and ubiquitin-Rhodamine (Ub-Rho) as its minimal fluorogenic substrate. The change in rhodamine fluorescence upon its cleavage from ubiquitin was used to follow the catalytic reaction. The enzymes were treated with increasing amounts of MS-8 ranging from 0 µM to 500 µM. The activity assays were conducted at 25°C in a buffer containing 20 mM Tris pH 8.0, 0.05% CHAPS, and 10 mM BME, in Corning black 384-well plates with a volume of 34 µL per well. DMSO served as a negative control. Fluorescence was measured every 11 seconds for 20 minutes using a Molecular Devices SpectraMax iD5 Multi-Mode Microplate Reader with excitation and emission wavelengths set to 487 nm and 535 nm, respectively. Protein concentrations were kept constant at 0.1 nM to hydrolyze Ub-Rho at a fixed substrate concentration of 500 nM.

A linear increase in raw fluorescence over time was used to obtain the initial velocity of the reaction (V_0_) at varying concentrations of MS-8. Their dependence on the concentration of MS-8 was analyzed in GraphPad Prism using the log(dose) vs. response equation with non-linear regression. All experiments were performed at least twice.

