## Supplementary Information for "Functional Spectrum of USP7 Pathogenic Variants in Hao-Fountain Syndrome: Insights into the Enzyme’s Activity, Stability, and Allosteric Modulation"

Figure S1

A

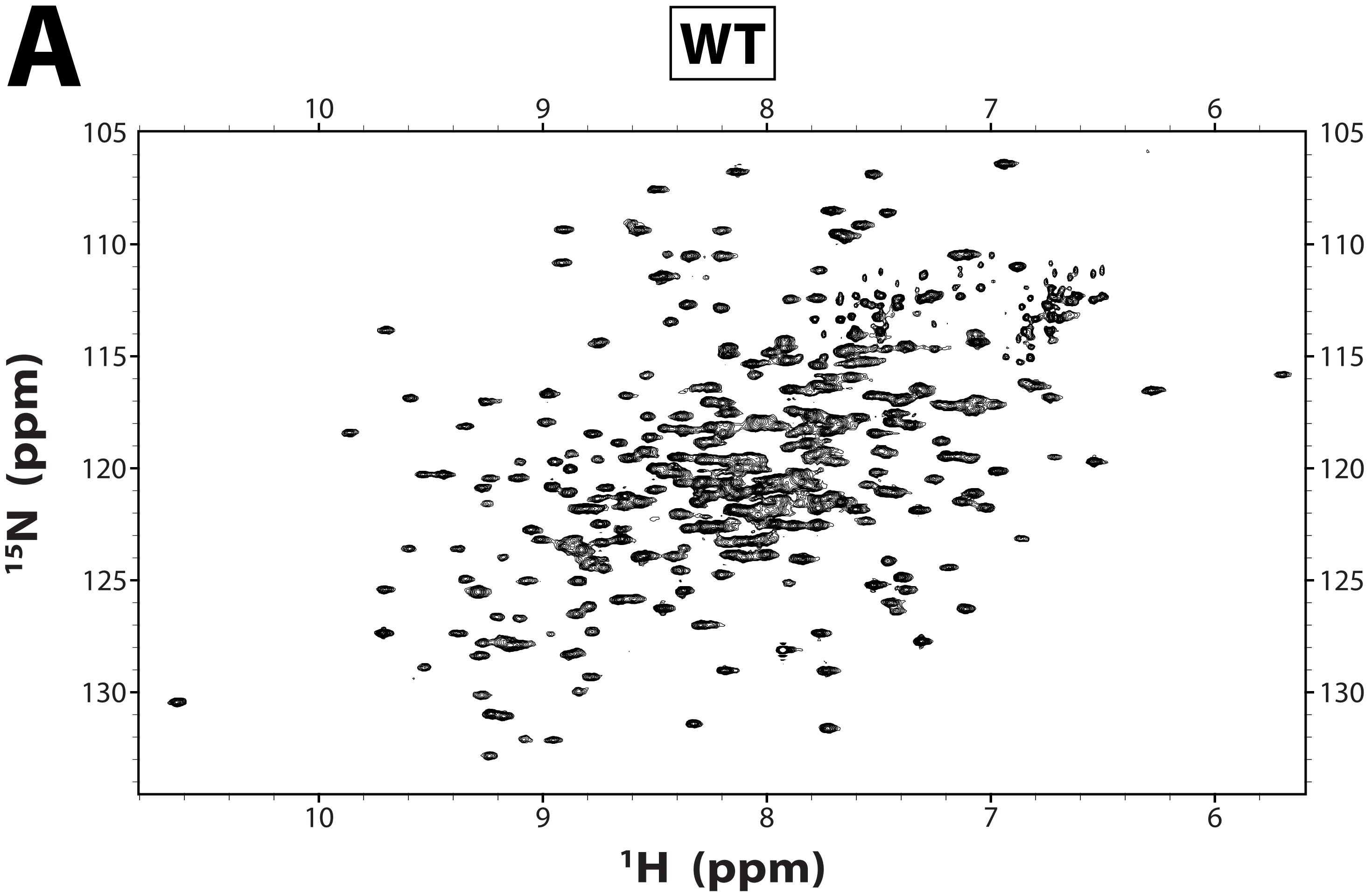

Figure S1

**B**

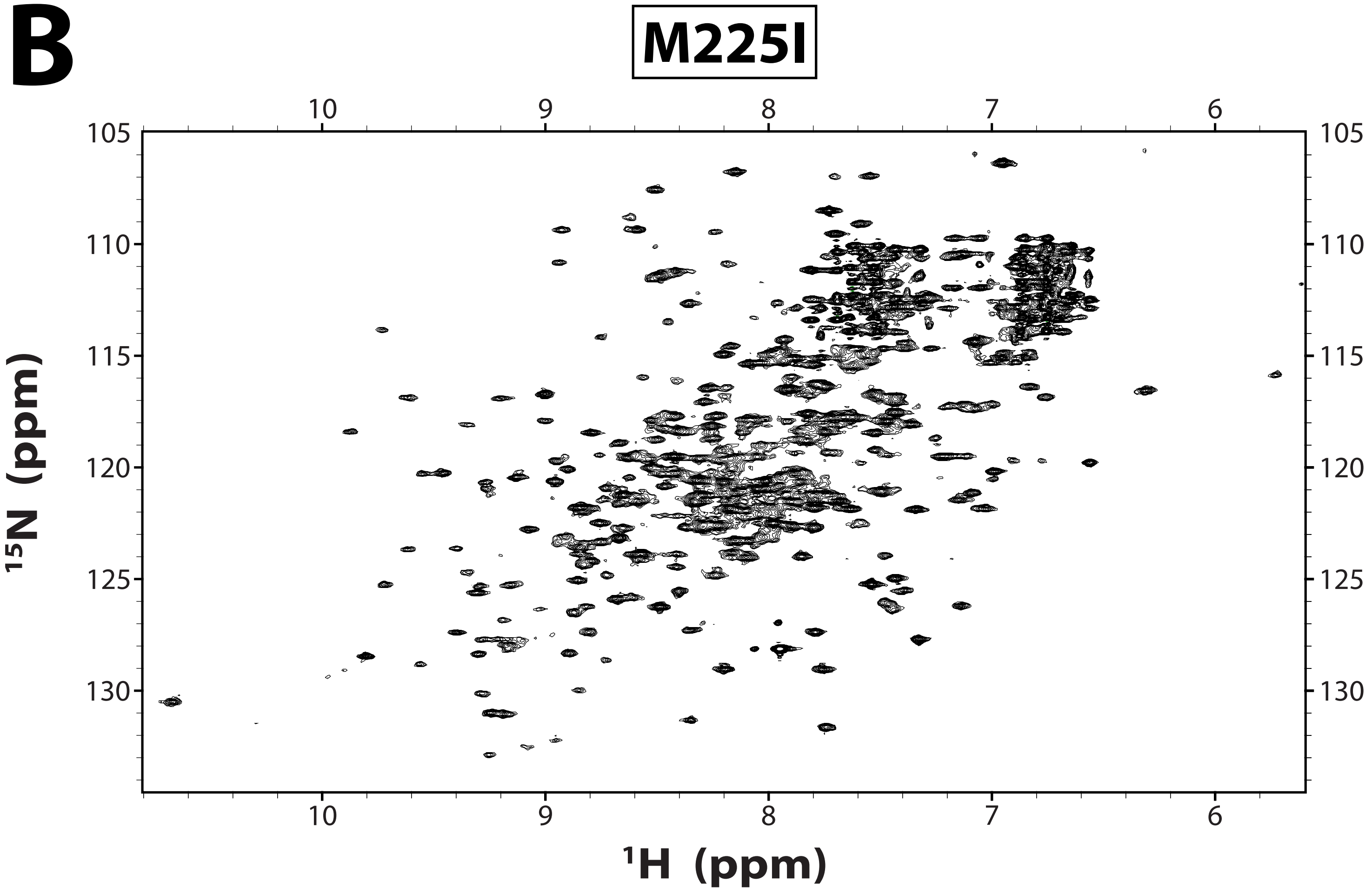

Figure S1

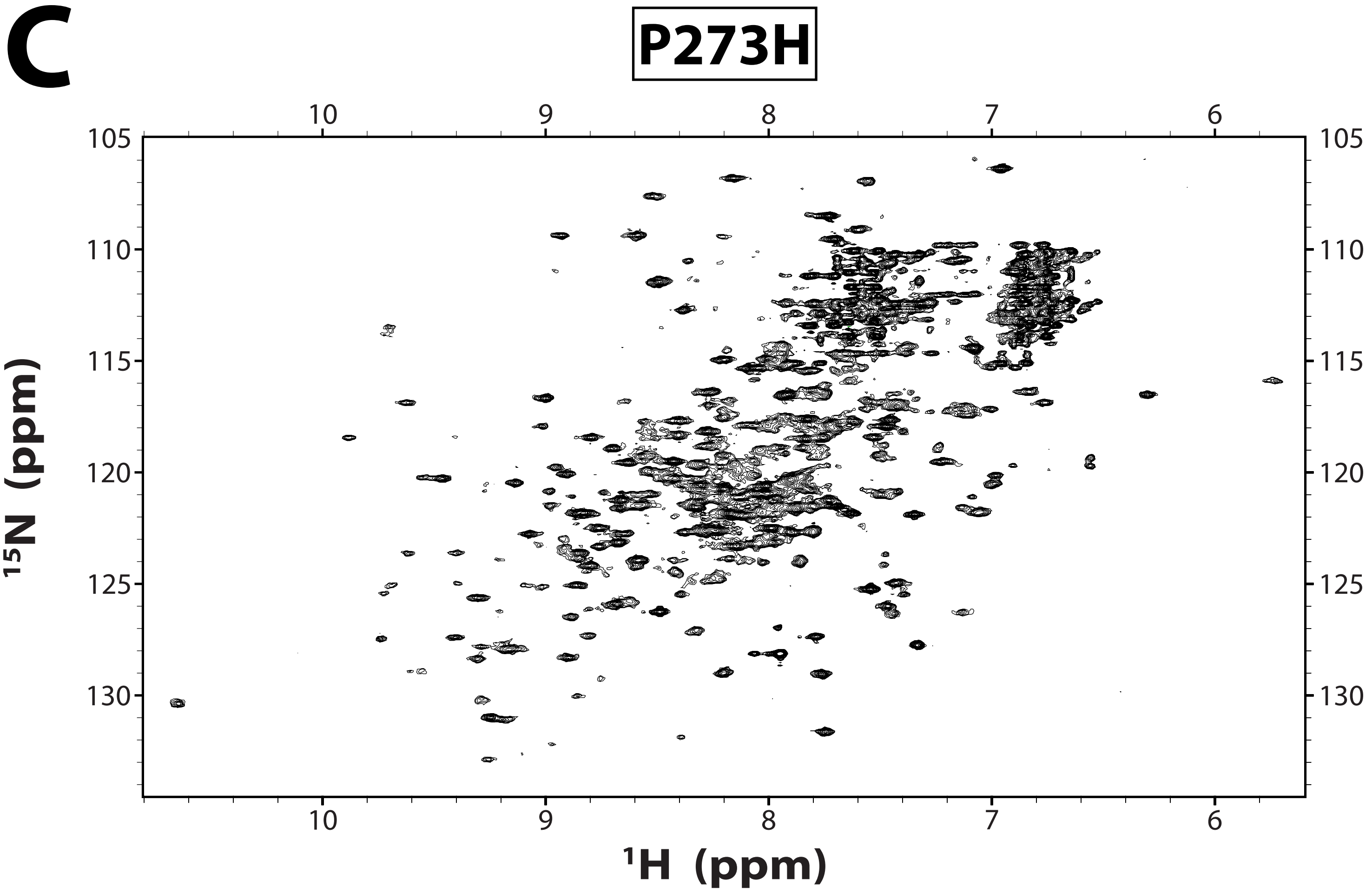

Figure S1

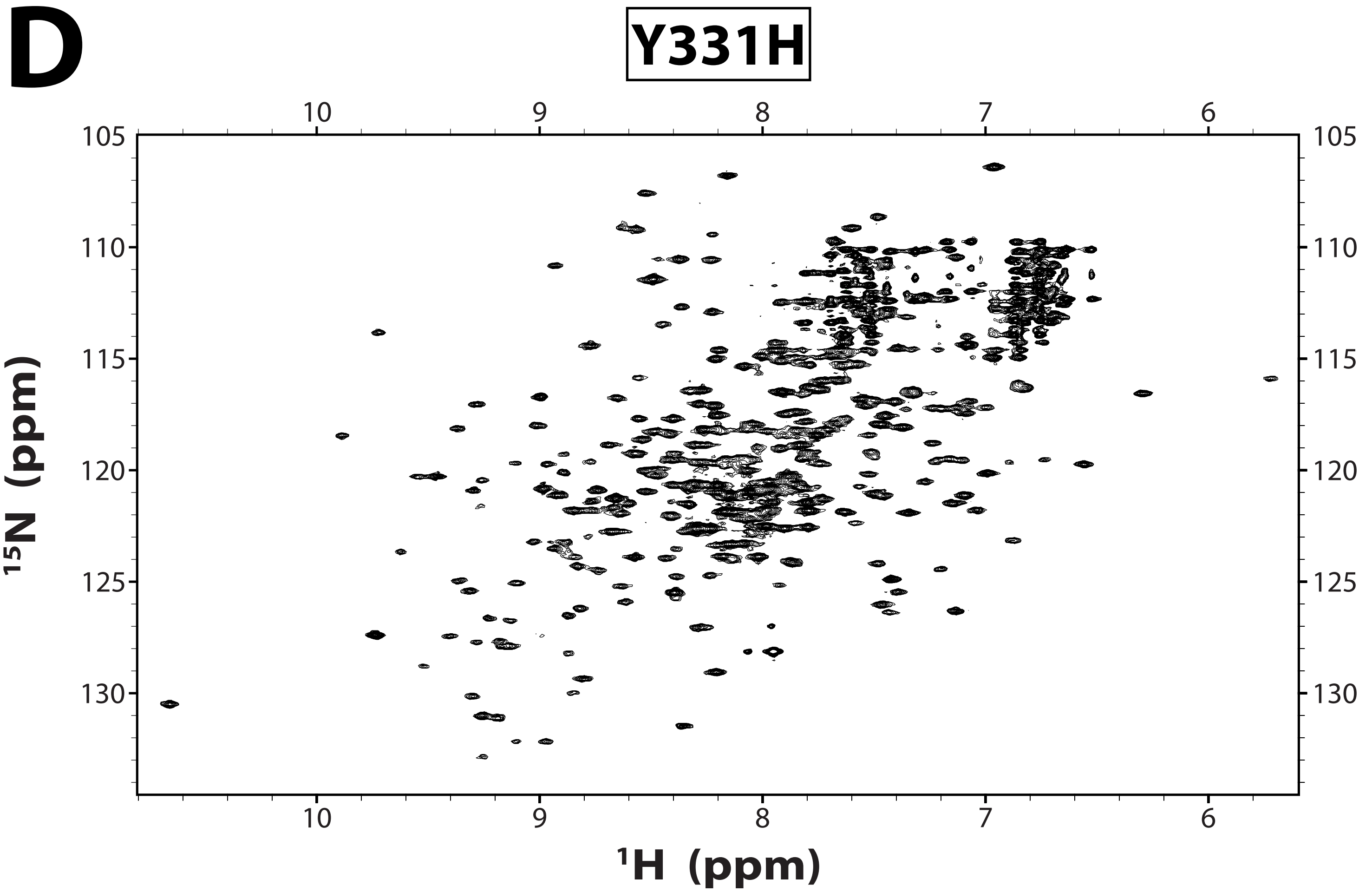

Figure S1

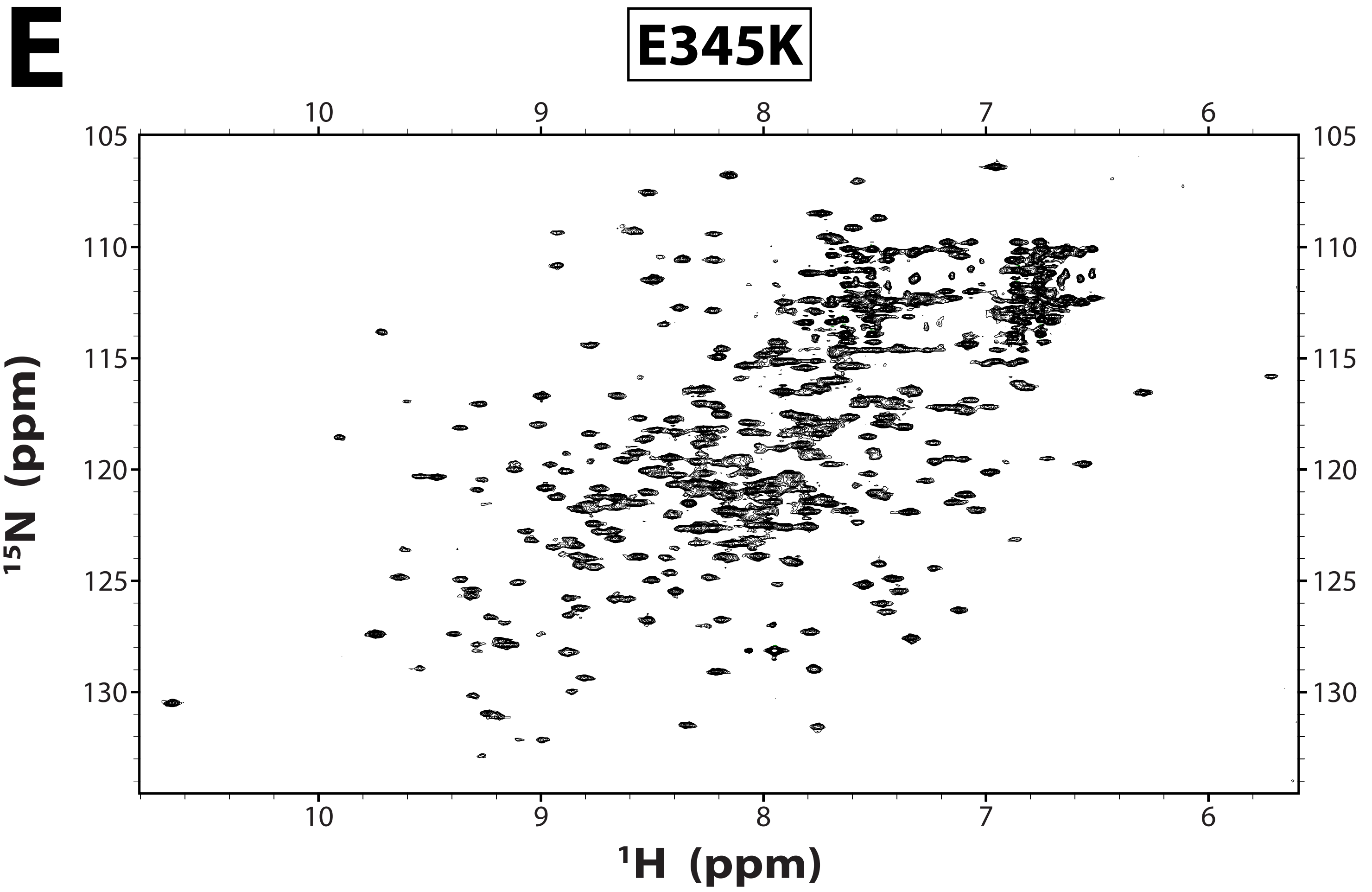

Figure S1

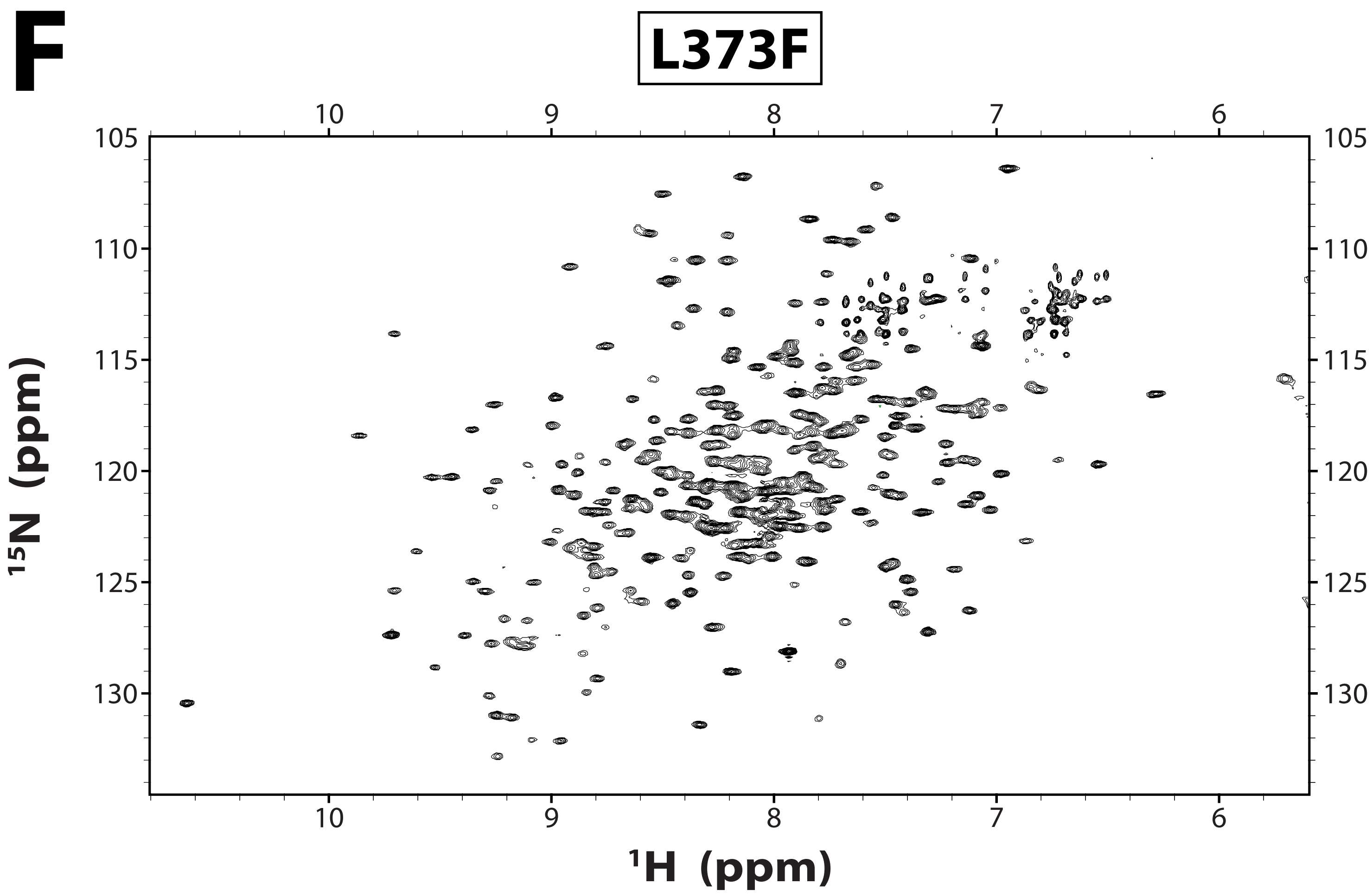

Figure S1

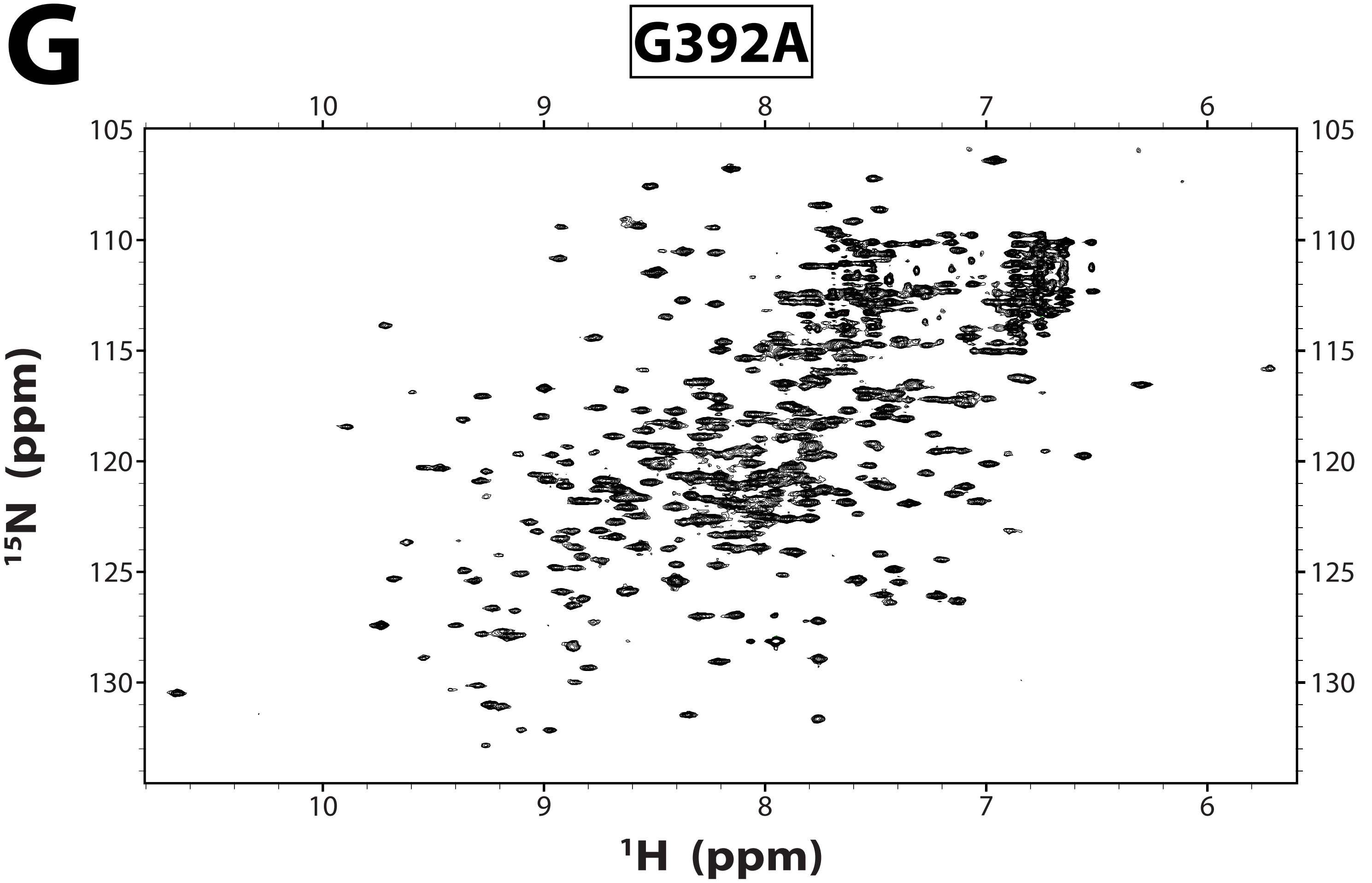

Figure S1

**H**

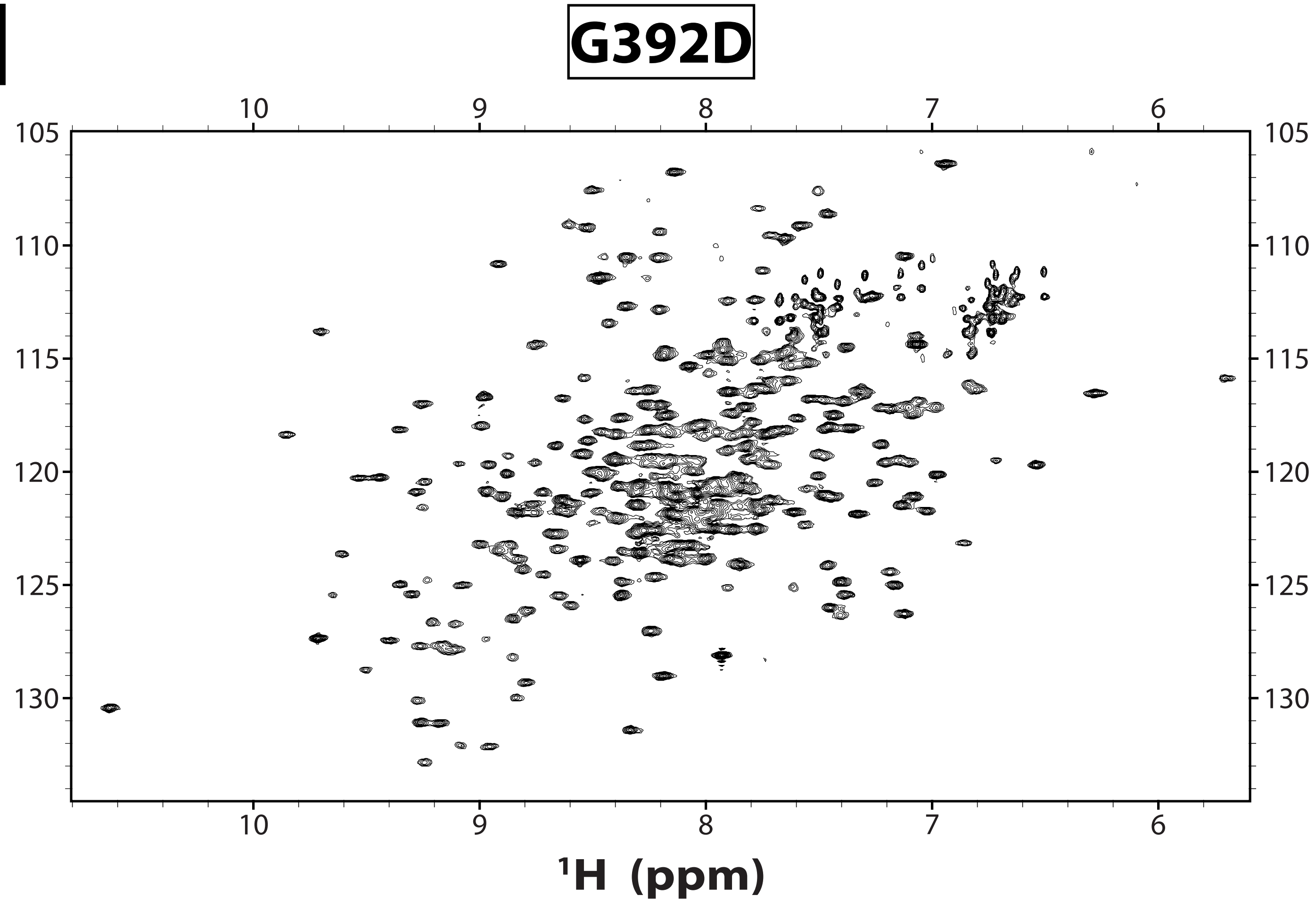

### Figure S1

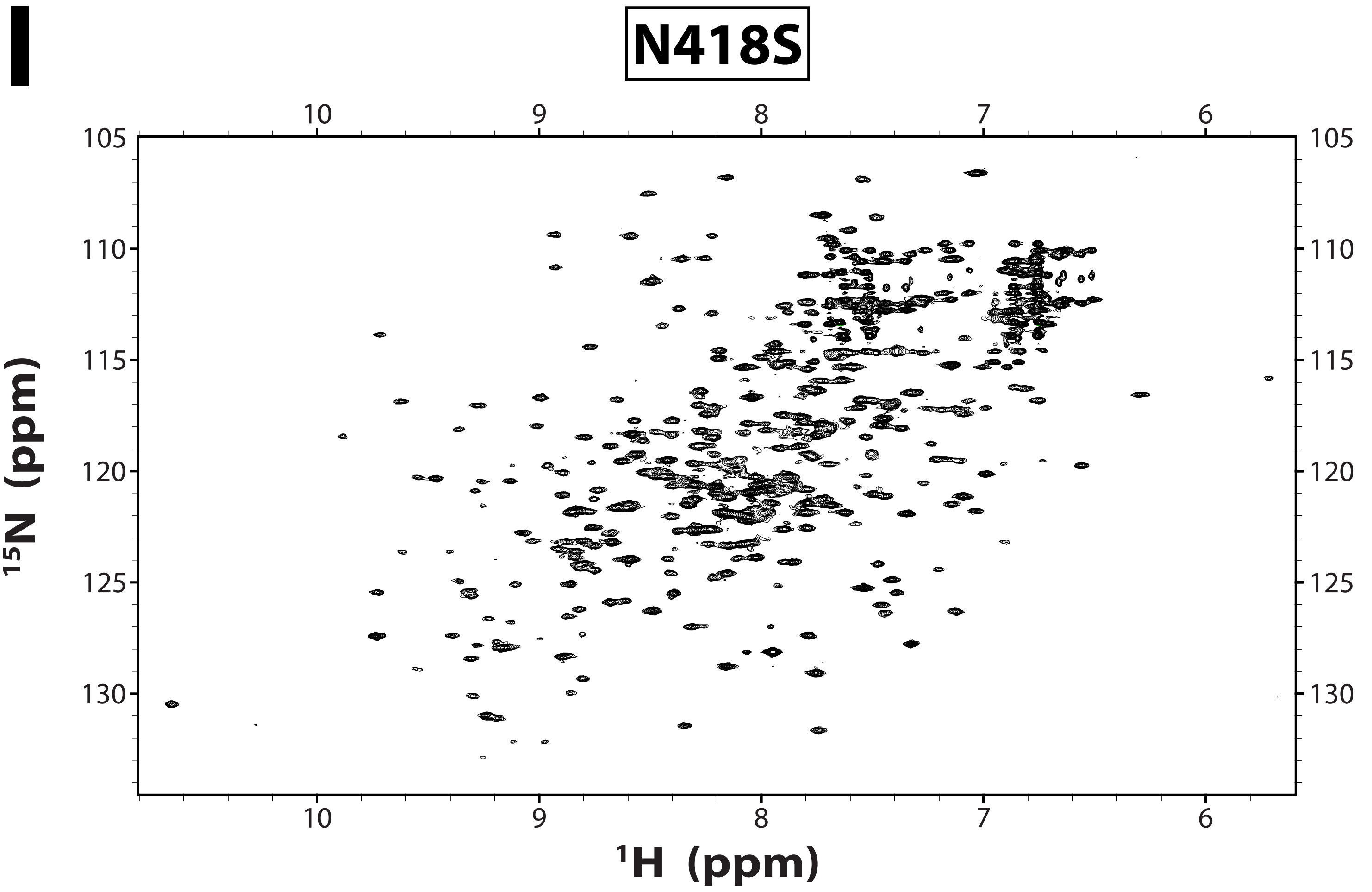

Figure S1

J

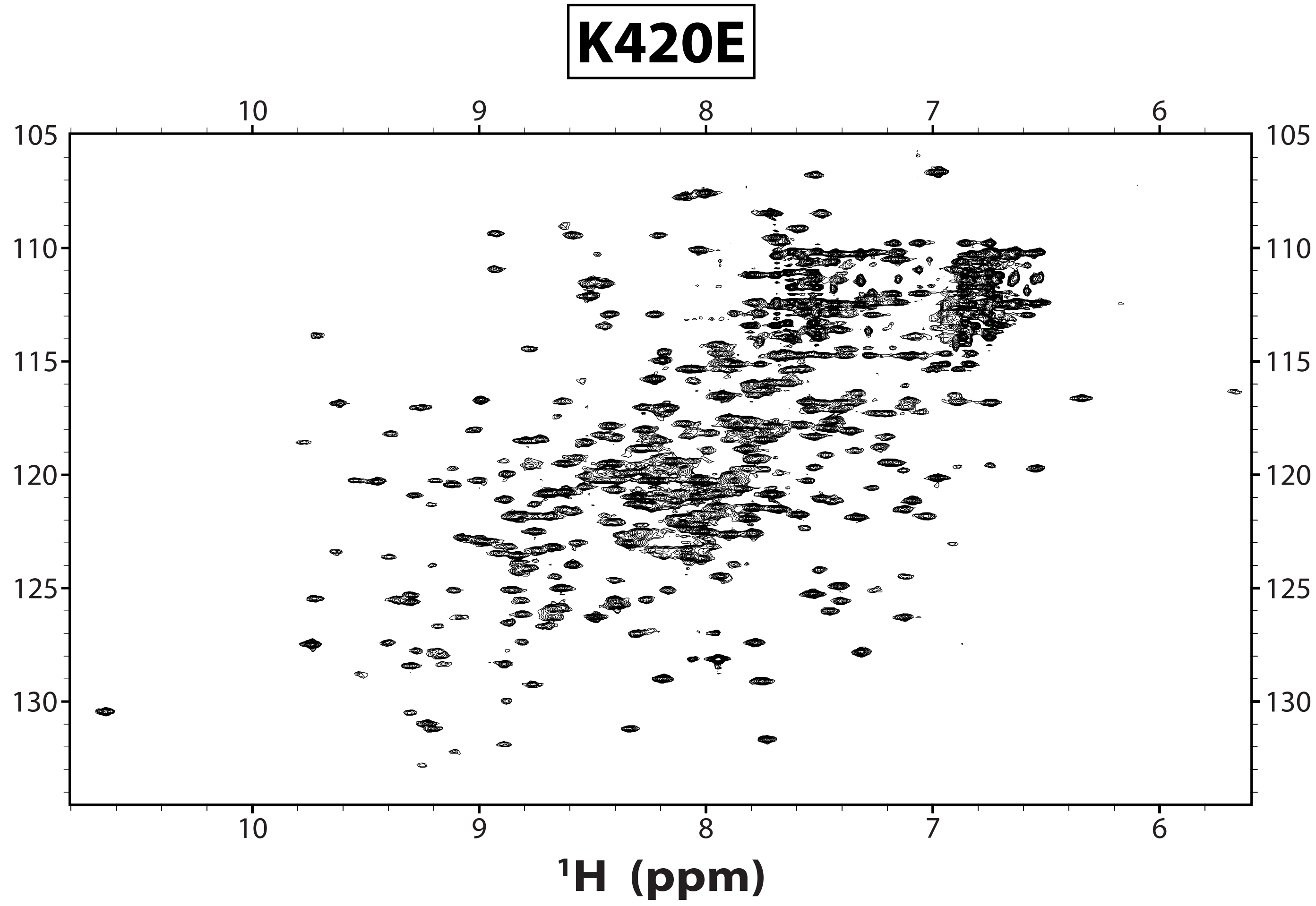

Figure S1

K

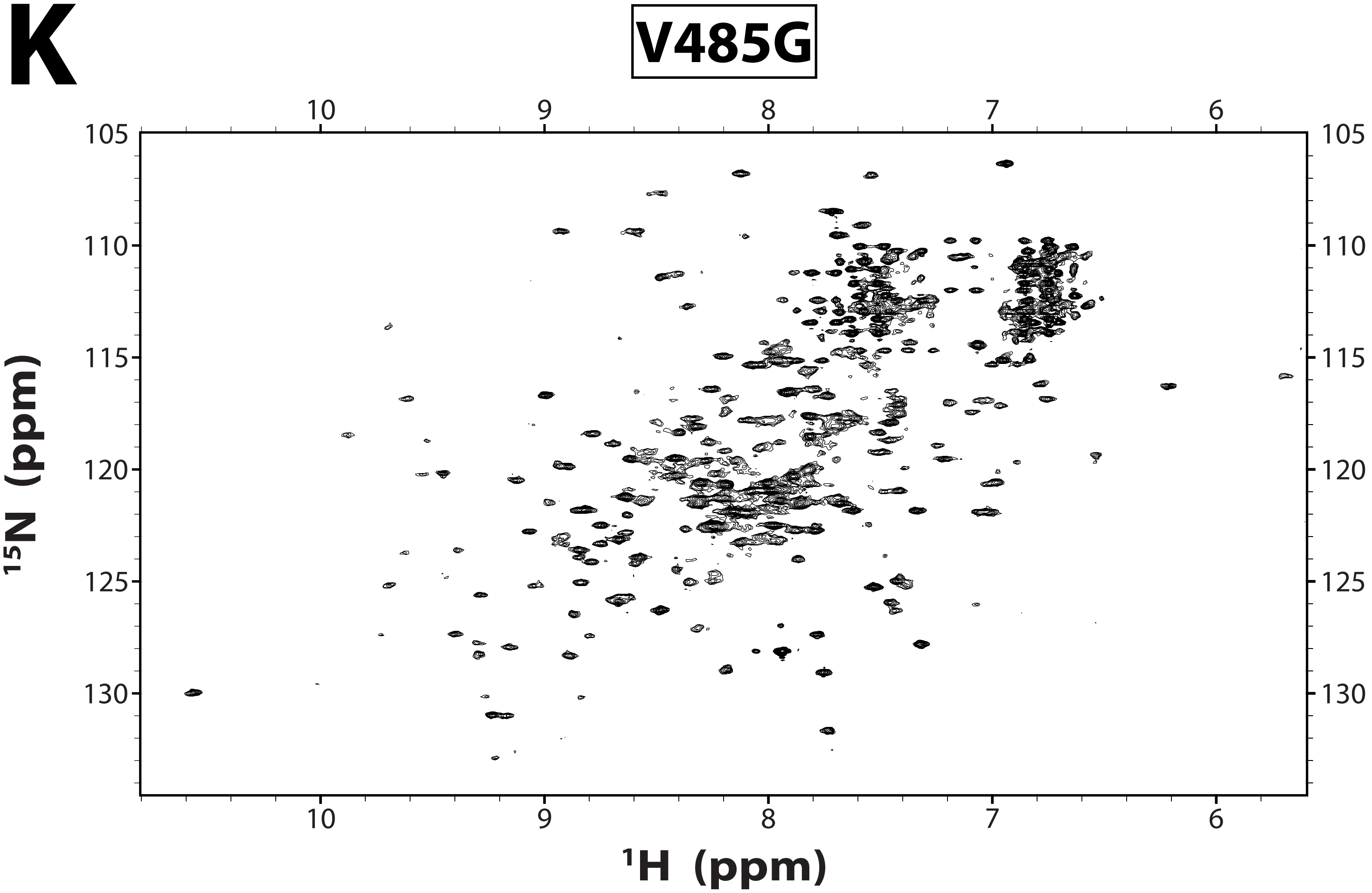

**Figure S1. NMR spectra of USP7 variants associated with Hao-Fountain syndrome.**
2D $^{15}N$ TROSY spectra of $^{15}N$-labeled USP7 catalytic domain and its Hao-Fountain syndrome variants. Individual spectra are labeled with the corresponding variant name.

### Figure S2

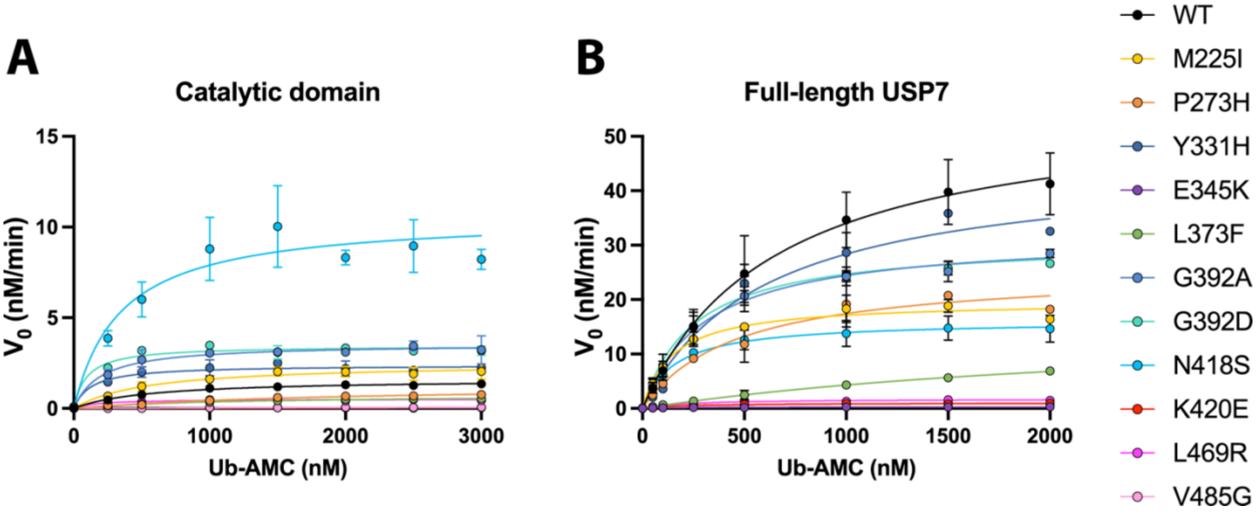


**Figure S2. Effect of Hao-Fountain syndrome variants on USP7 activity.**
Michaelis-Menten plots of the initial velocity ($V_0$) as a function of ubiquitin-AMC concentration shown for (**A**) USP7 catalytic domain and its mutants and (**B)** FL-USP7 and its mutants.

### Figure S3

## A

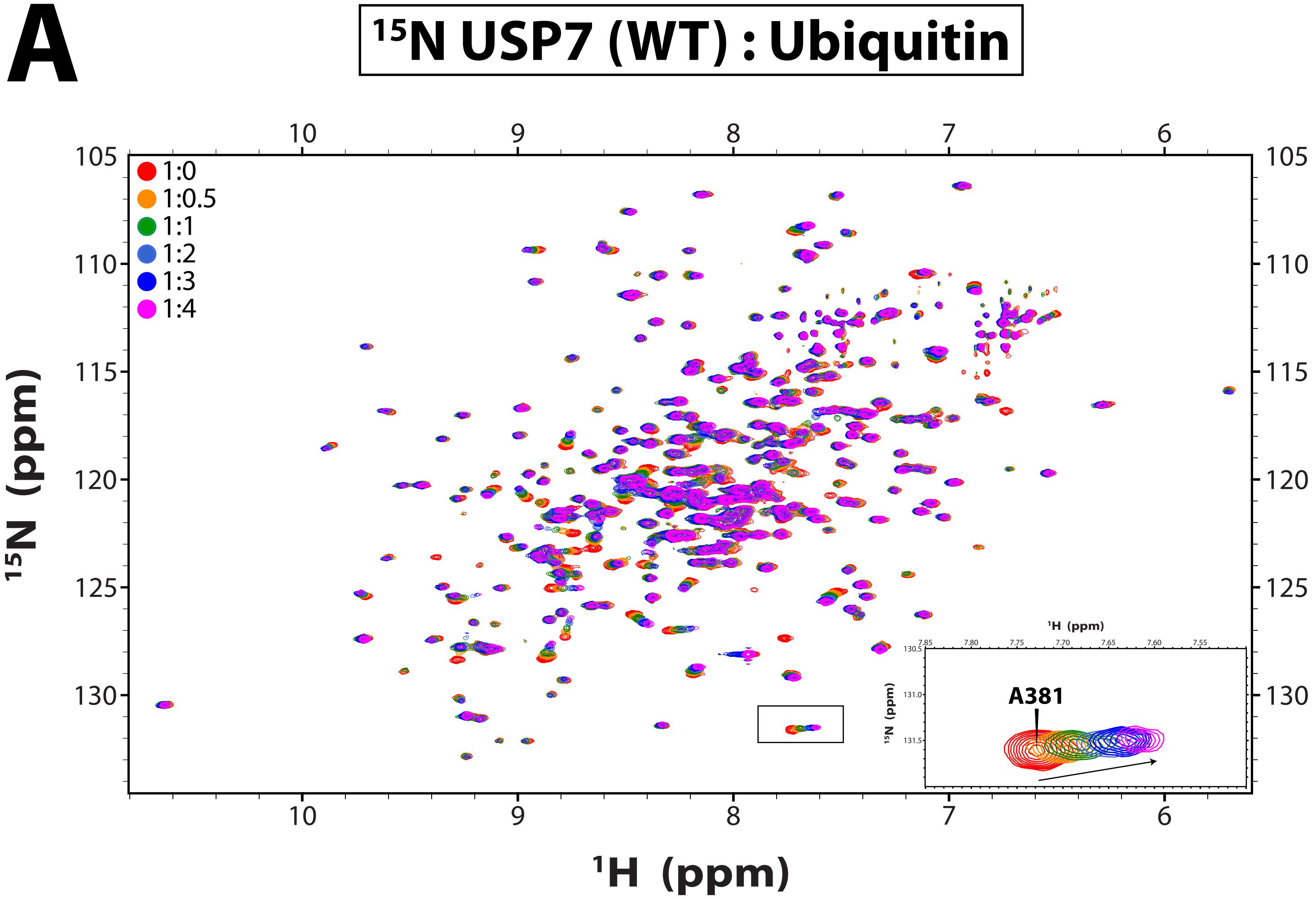

15N USP7 (WT) : Ubiquitin
1:0
1:0.5
1:1
1:2
1:3
1:4
15N (ppm)
1H (ppm)
A381


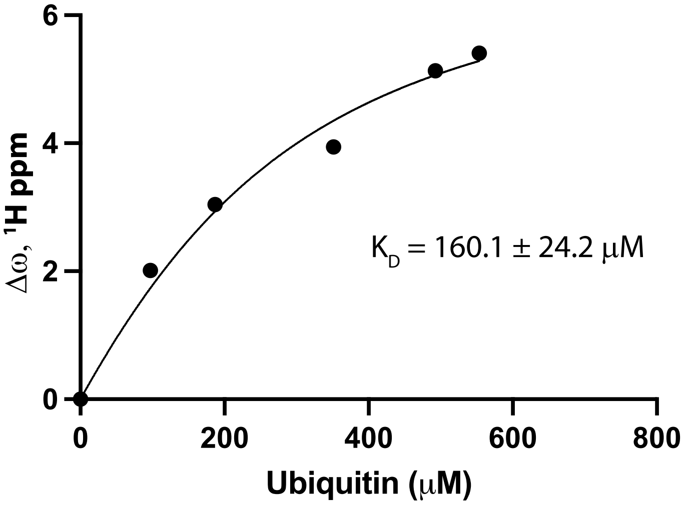

Δω, 1H ppm
K_D = 160.1 ± 24.2 μM
Ubiquitin (μM)

Figure S3

**B**

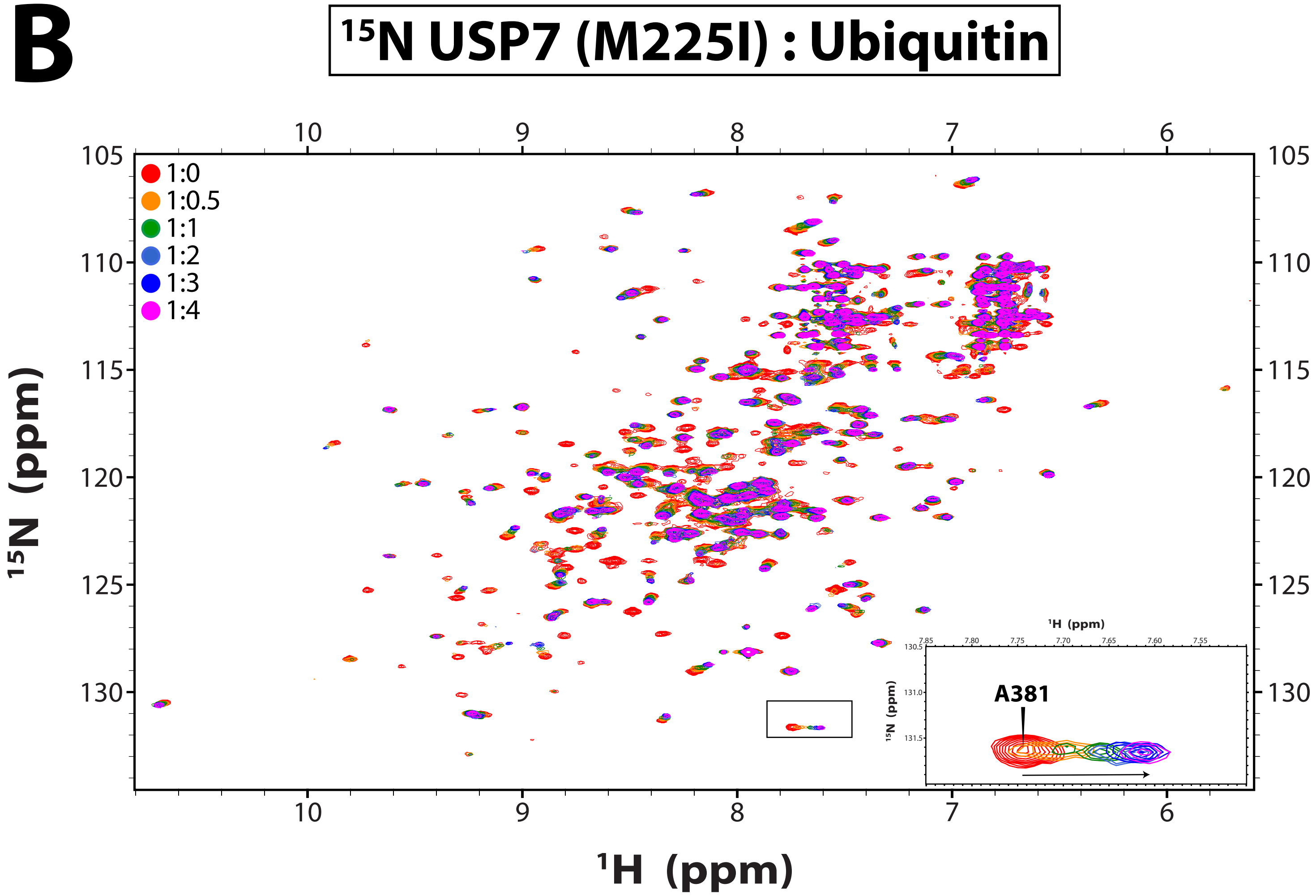

$^{15}$N USP7 (M225I) : Ubiquitin
1:0
1:0.5
1:1
1:2
1:3
1:4
A381
$^{15}$N (ppm)
$^{1}$H (ppm)


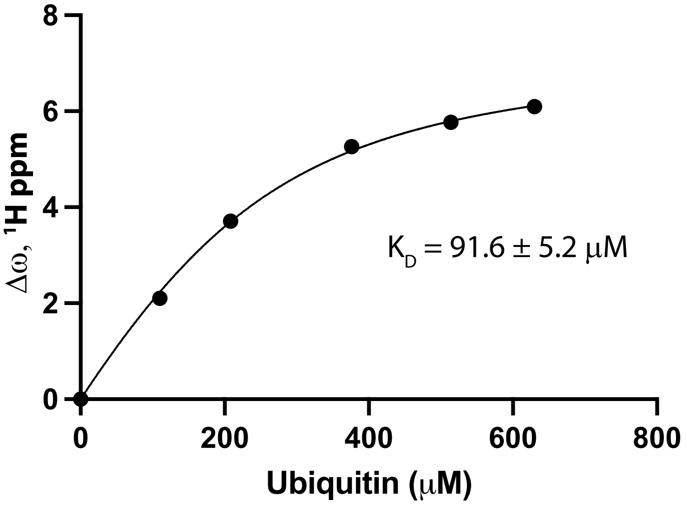

K$_D$ = 91.6 ± 5.2 μM
Δω, $^{1}$H ppm
Ubiquitin (μM)

Figure S3

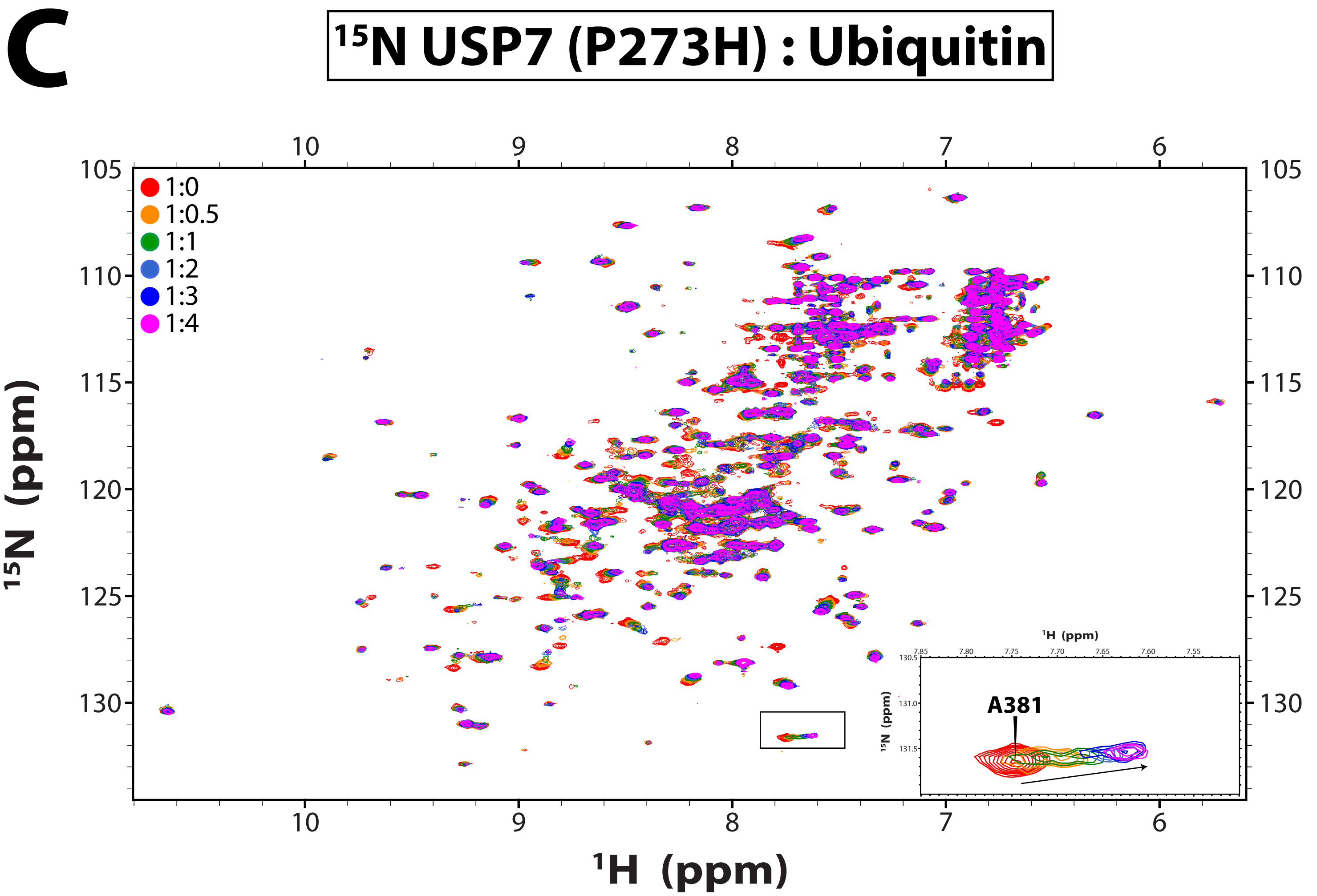
C
$^{15}$N USP7 (P273H) : Ubiquitin
1:0
1:0.5
1:1
1:2
1:3
1:4
A381
$^{15}$N (ppm)
$^{1}$H (ppm)

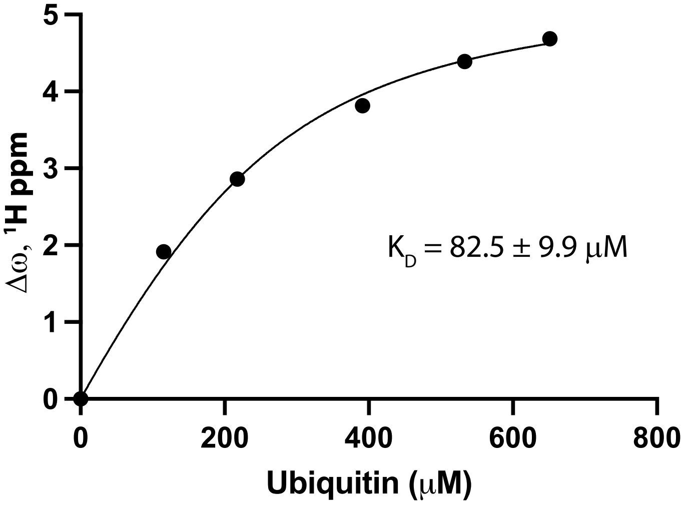
$K_D$ = 82.5 ± 9.9 μM
Δω, $^{1}$H ppm
Ubiquitin (μM)

### Figure S3

**D**

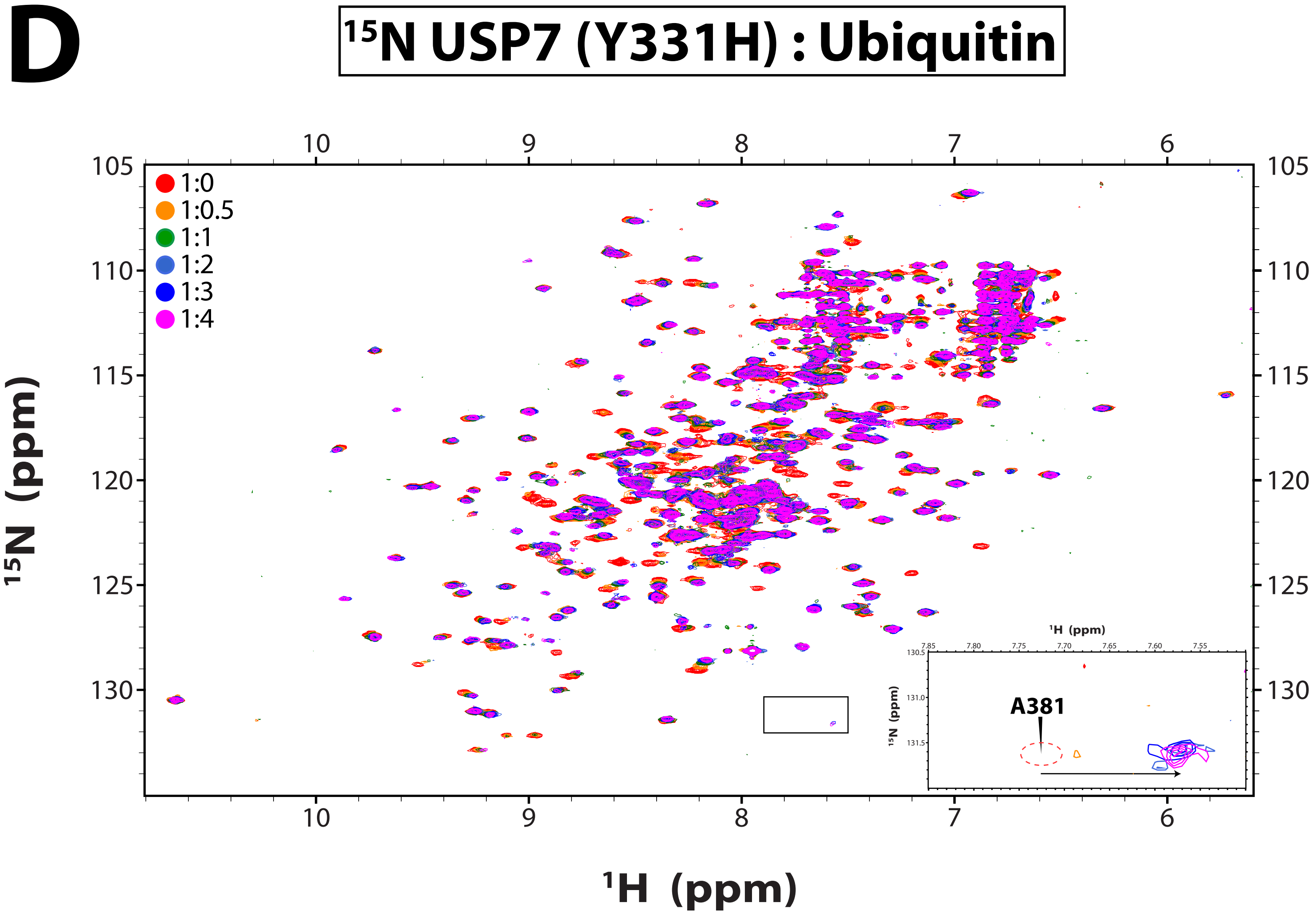

¹⁵N USP7 (Y331H) : Ubiquitin
1:0
1:0.5
1:1
1:2
1:3
1:4
¹⁵N (ppm)
¹H (ppm)
10
9
8
7
6
105
110
115
120
125
130
A381
7.85
7.80
7.75
7.70
7.65
7.60
7.55
130.5
131.0
131.5


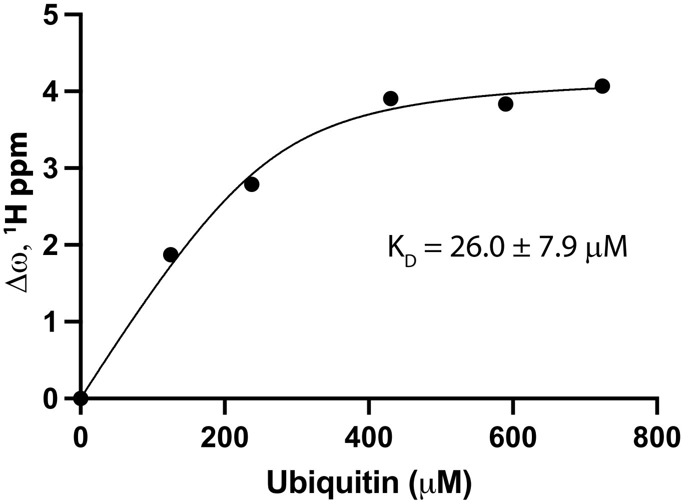

Δω, ¹H ppm
Ubiquitin (μM)
K_D = 26.0 ± 7.9 μM
0
1
2
3
4
5
200
400
600
800

Figure S3

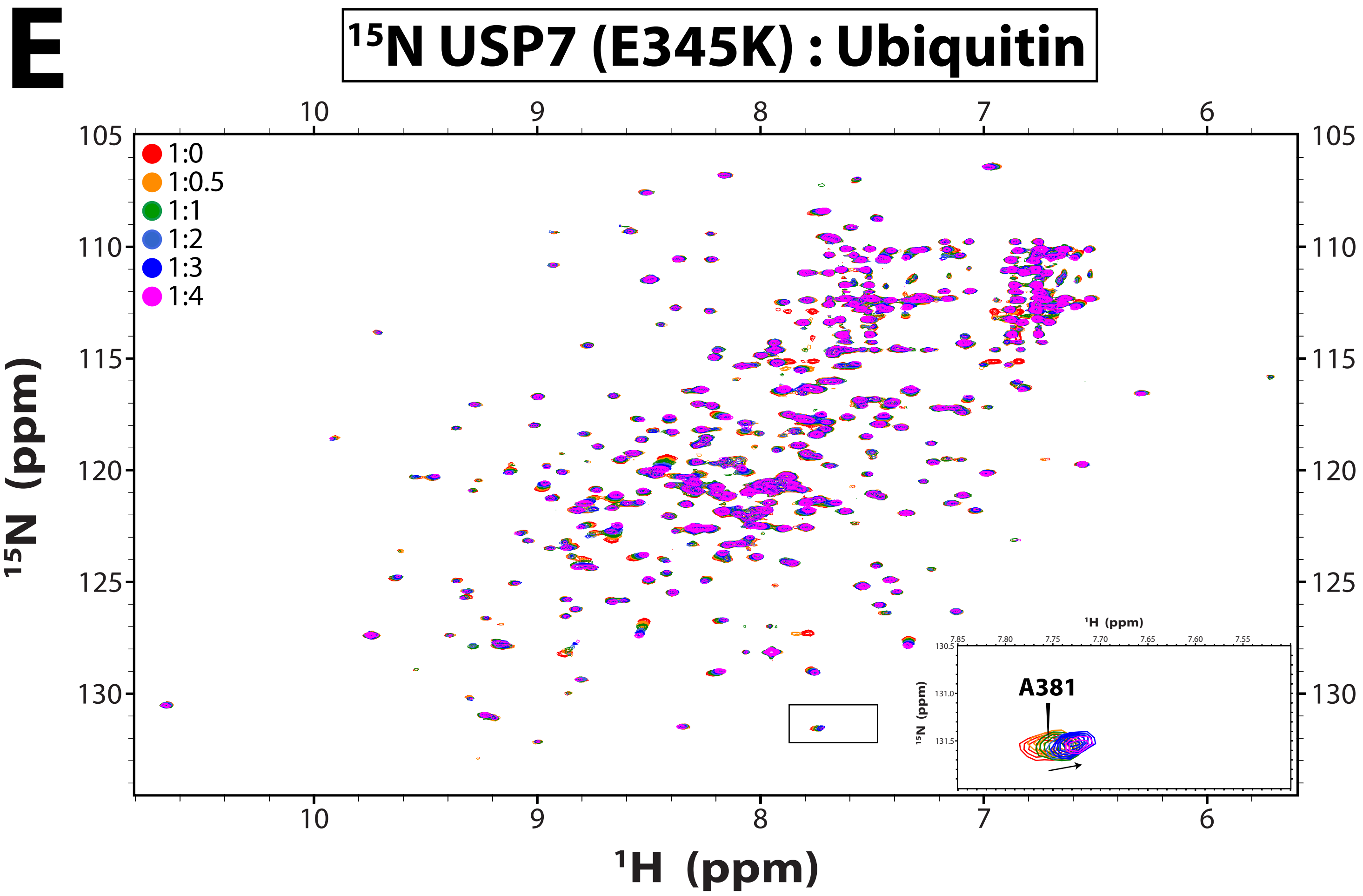
E
15N USP7 (E345K) : Ubiquitin
1:0
1:0.5
1:1
1:2
1:3
1:4
A381
1H (ppm)
15N (ppm)

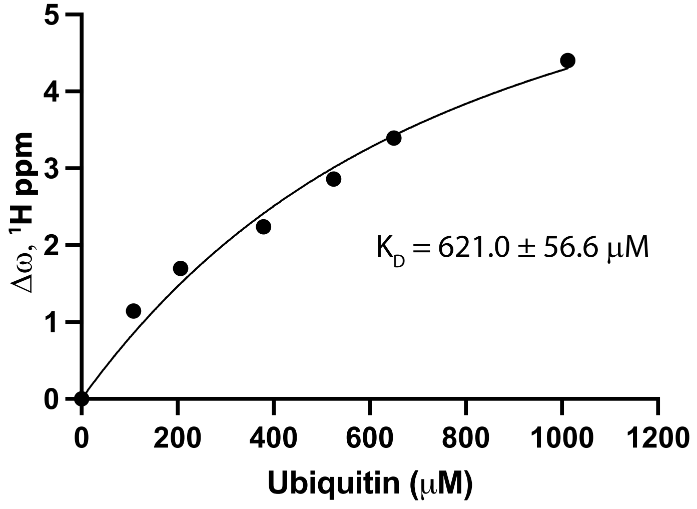
$K_D$ = 621.0 ± 56.6 μM
Δω, 1H ppm
Ubiquitin (μM)

Figure S3

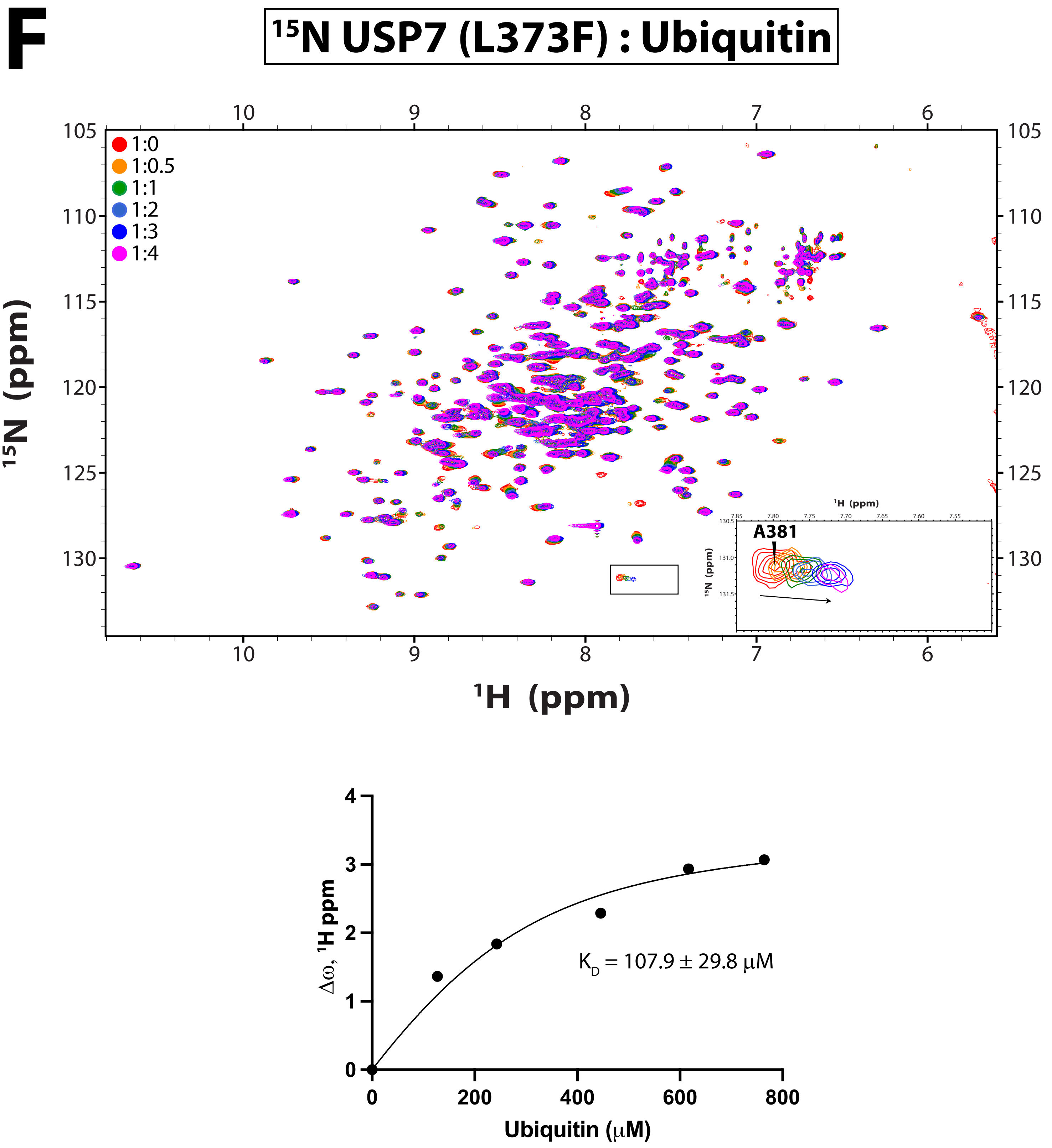
F
15N USP7 (L373F) : Ubiquitin
1:0
1:0.5
1:1
1:2
1:3
1:4
15N (ppm)
1H (ppm)
A381
Δω, 1H ppm
Ubiquitin (μM)
KD = 107.9 ± 29.8 μM

### Figure S3

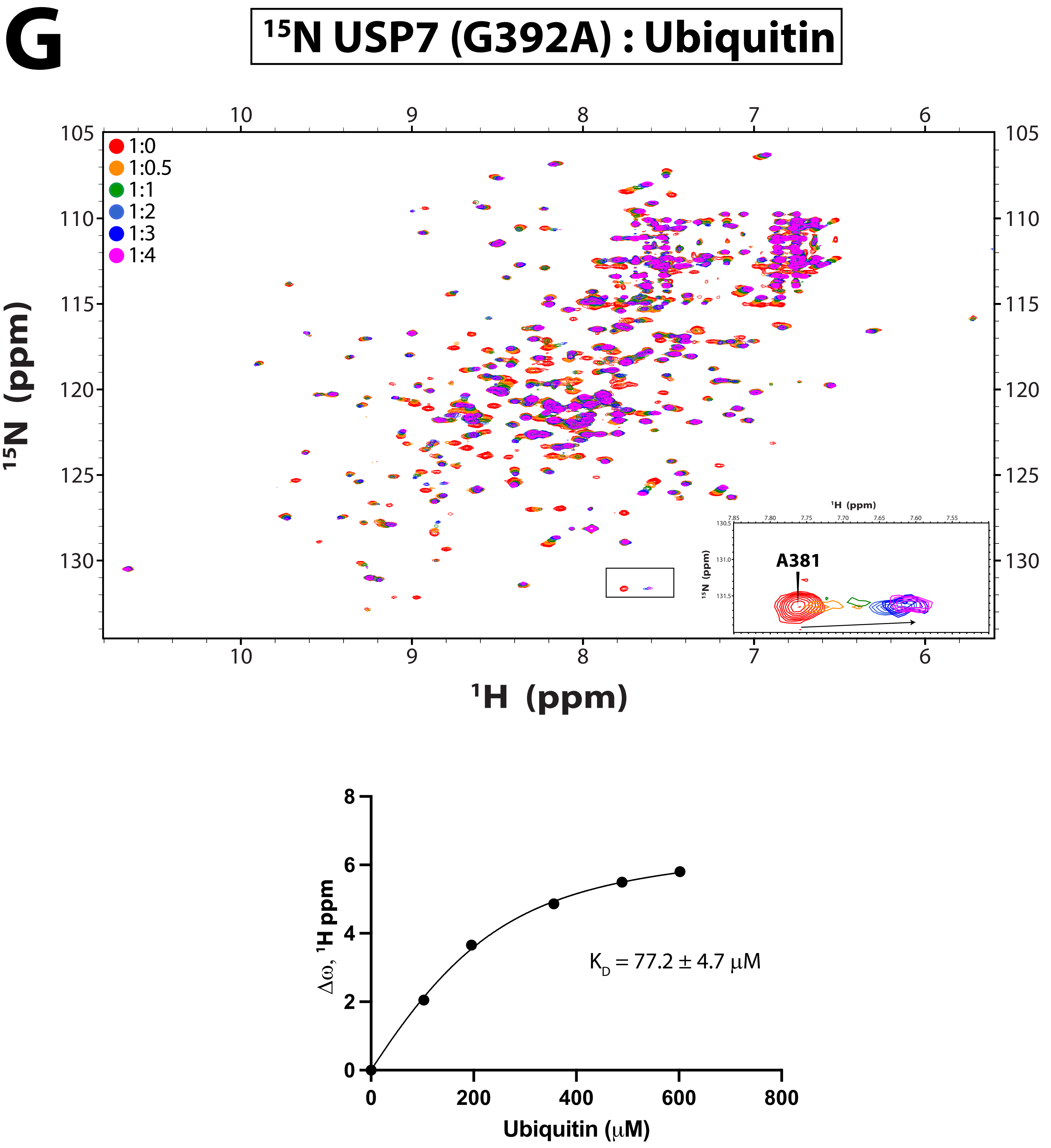
G
15N USP7 (G392A) : Ubiquitin
1:0
1:0.5
1:1
1:2
1:3
1:4
15N (ppm)
1H (ppm)
A381
KD = 77.2 ± 4.7 μM
Δω, 1H ppm
Ubiquitin (μM)

### Figure S3

# H

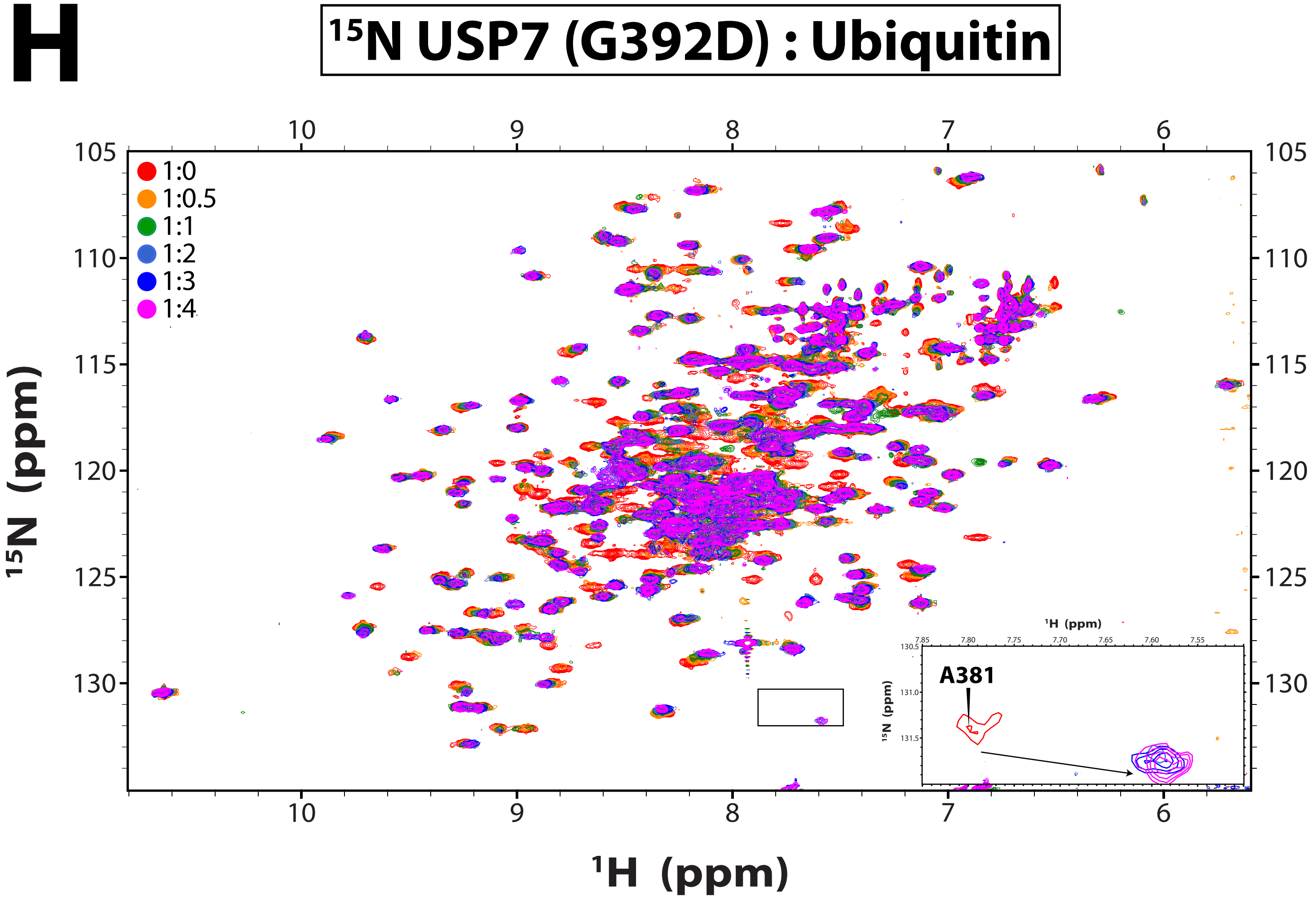

15N USP7 (G392D) : Ubiquitin
1:0
1:0.5
1:1
1:2
1:3
1:4
A381
15N (ppm)
1H (ppm)


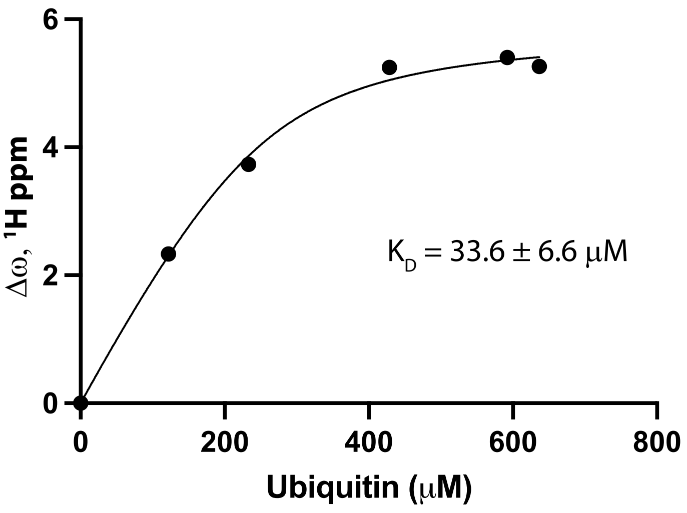

KD = 33.6 ± 6.6 μM
Δω, 1H ppm
Ubiquitin (μM)

### Figure S3

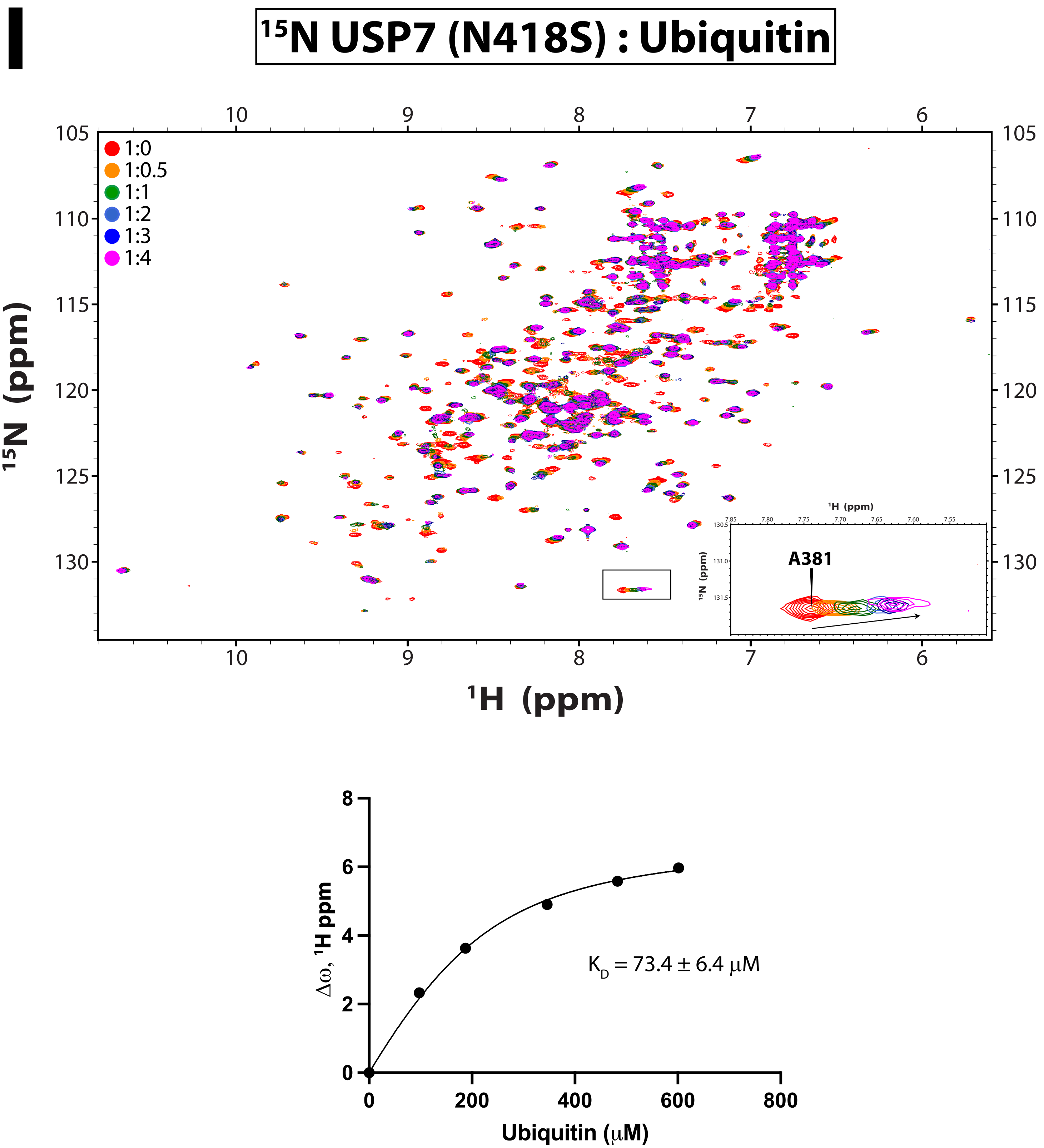
I
15N USP7 (N418S) : Ubiquitin
1:0
1:0.5
1:1
1:2
1:3
1:4
15N (ppm)
1H (ppm)
A381
KD = 73.4 ± 6.4 μM
Δω, 1H ppm
Ubiquitin (μM)

Figure S3

**J**

**$^{15}$N USP7 (K420E) : Ubiquitin**

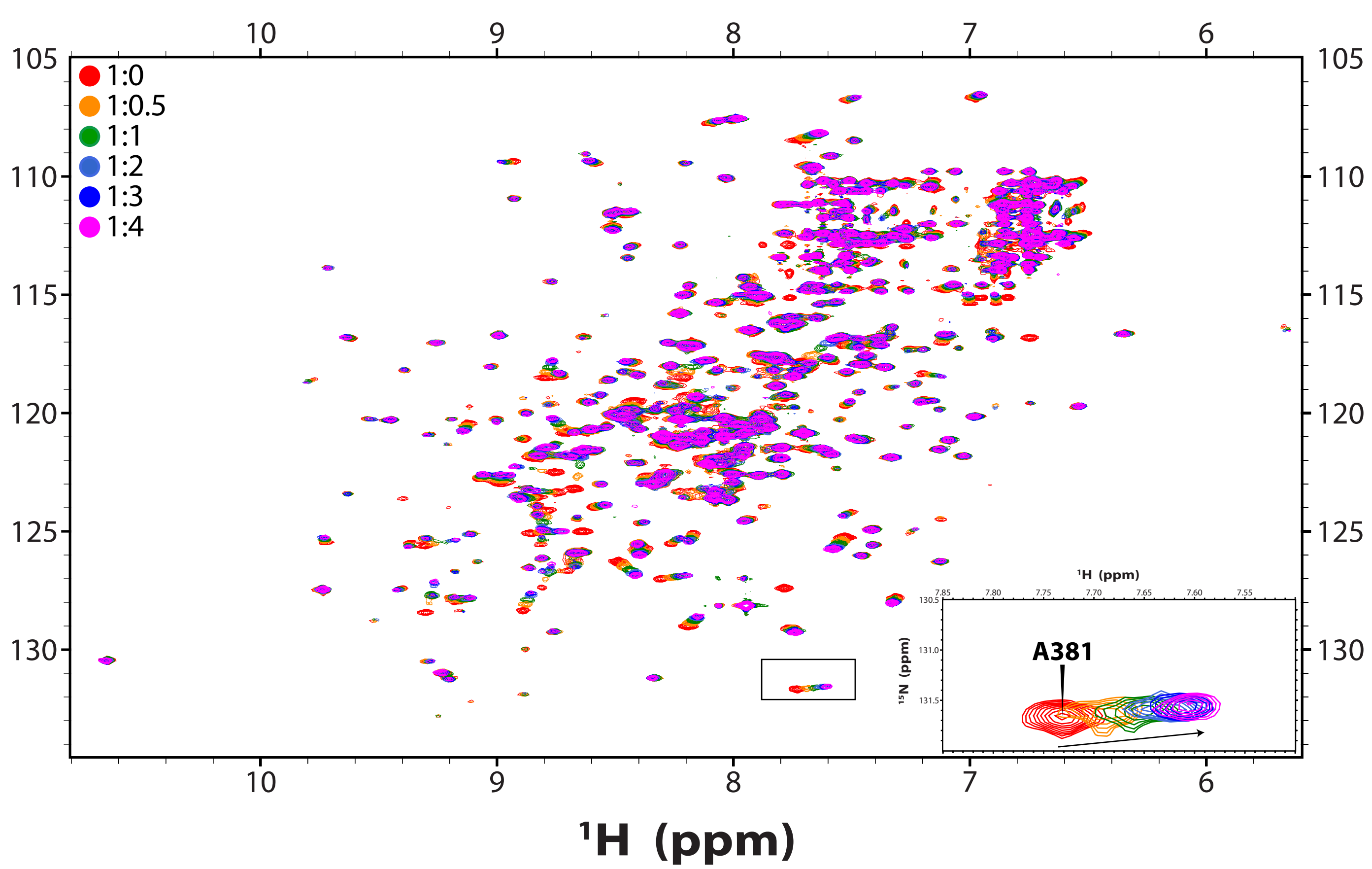


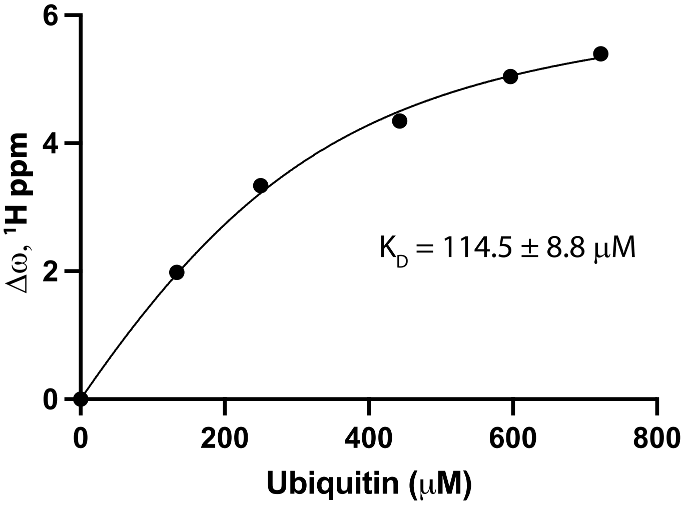

### Figure S3

**K**

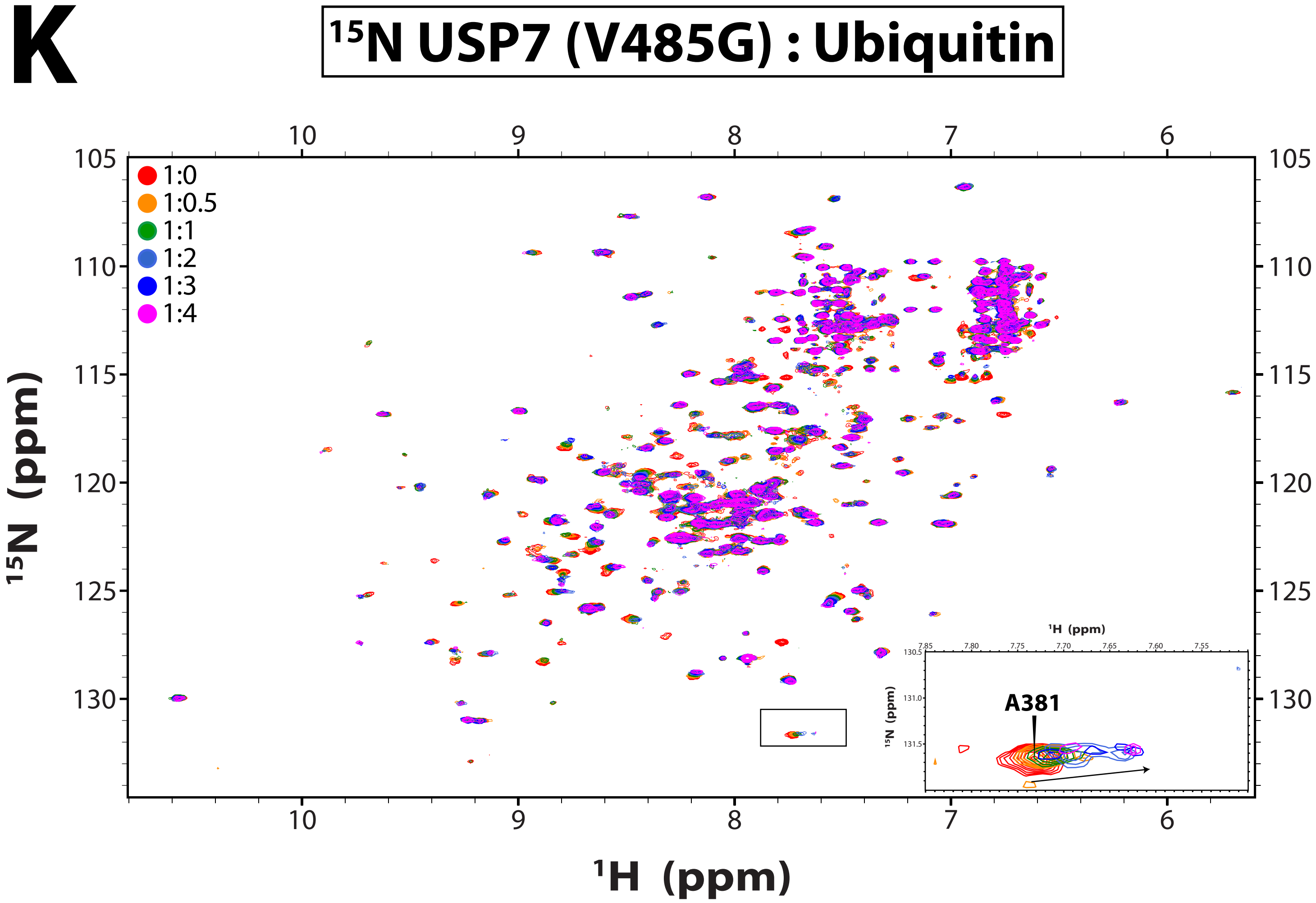

15N USP7 (V485G) : Ubiquitin
1:0
1:0.5
1:1
1:2
1:3
1:4
A381
15N (ppm)
1H (ppm)


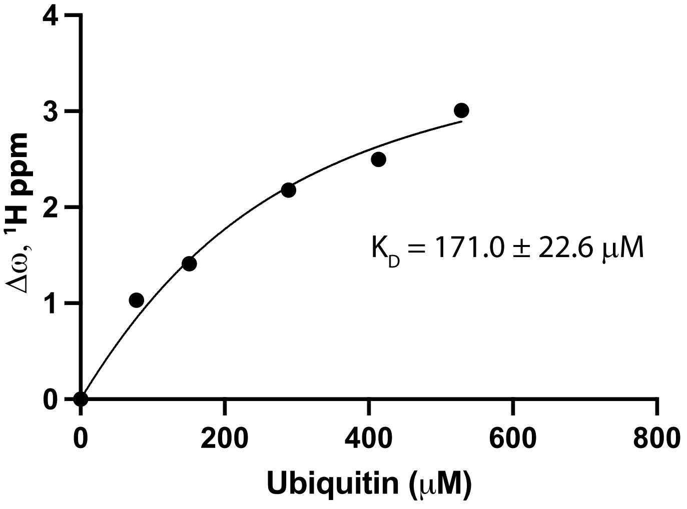

Δω, 1H ppm
Ubiquitin (μM)
K_D = 171.0 ± 22.6 μM

**Figure S3. Ubiquitin binding to catalytic domains of USP7.**

**Top:** $^{15}$N TROSY spectra of the $^{15}$N-labeled USP7 catalytic domain and its mutants gradually titrated with unlabeled ubiquitin. Residue A381 is showcased for each spectrum. USP7:ubiquitin molar ratios are shown. **Bottom:** Plot showing the global chemical shift perturbations ($\Delta\omega$) in the spectra as a function of ubiquitin concentration, used to estimate the binding affinities for each USP7 variant ($K_D$).

### Figure S4

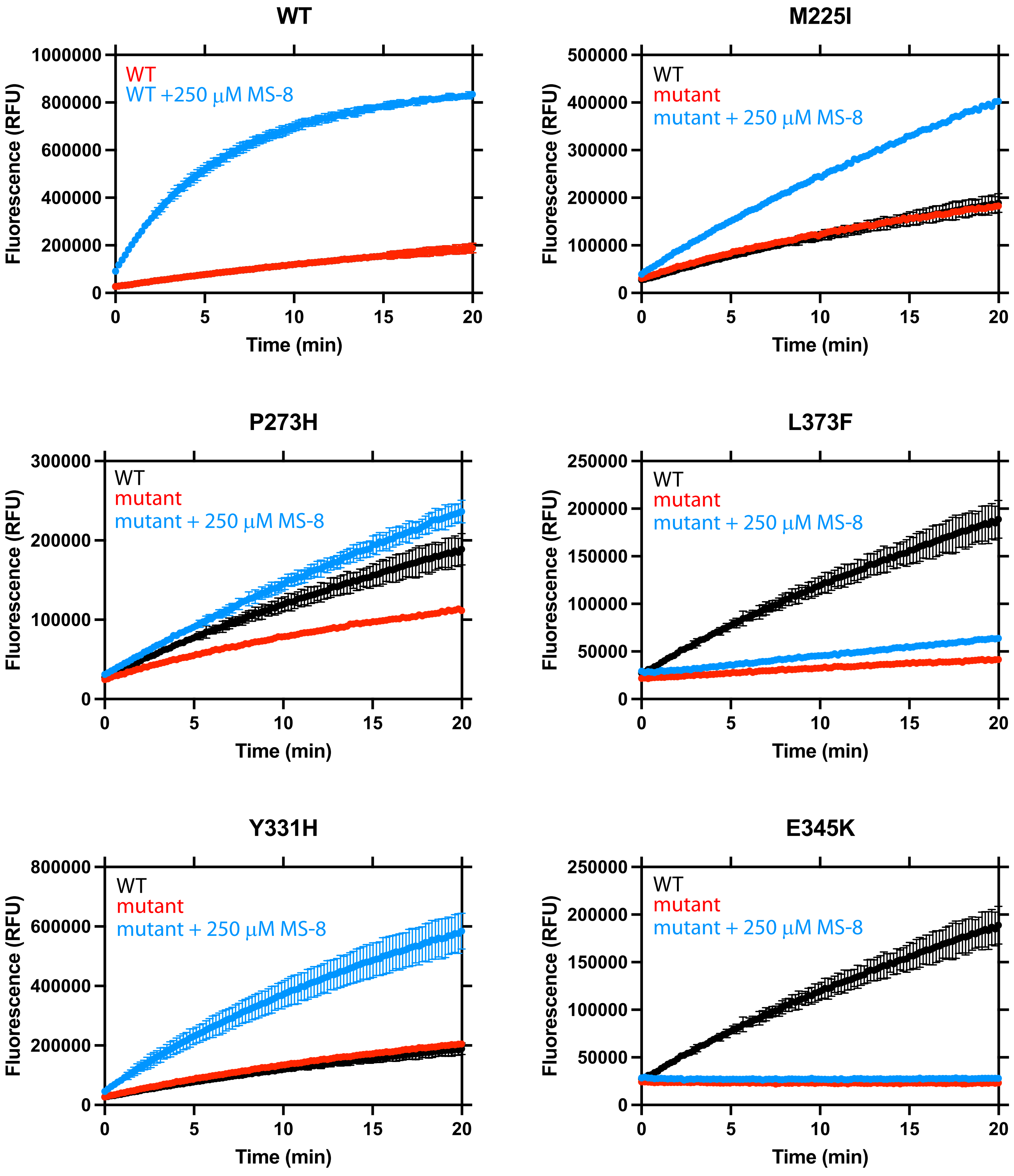
WT
1000000
800000
600000
400000
200000
0
WT
WT +250 µM MS-8
Fluorescence (RFU)
0
5
10
15
20
Time (min)
M225I
500000
400000
300000
200000
100000
0
WT
mutant
mutant + 250 µM MS-8
Fluorescence (RFU)
0
5
10
15
20
Time (min)
P273H
300000
200000
100000
0
WT
mutant
mutant + 250 µM MS-8
Fluorescence (RFU)
0
5
10
15
20
Time (min)
L373F
250000
200000
150000
100000
50000
0
WT
mutant
mutant + 250 µM MS-8
Fluorescence (RFU)
0
5
10
15
20
Time (min)
Y331H
800000
600000
400000
200000
0
WT
mutant
mutant + 250 µM MS-8
Fluorescence (RFU)
0
5
10
15
20
Time (min)
E345K
250000
200000
150000
100000
50000
0
WT
mutant
mutant + 250 µM MS-8
Fluorescence (RFU)
0
5
10
15
20
Time (min)

### Figure S4 (cont.)

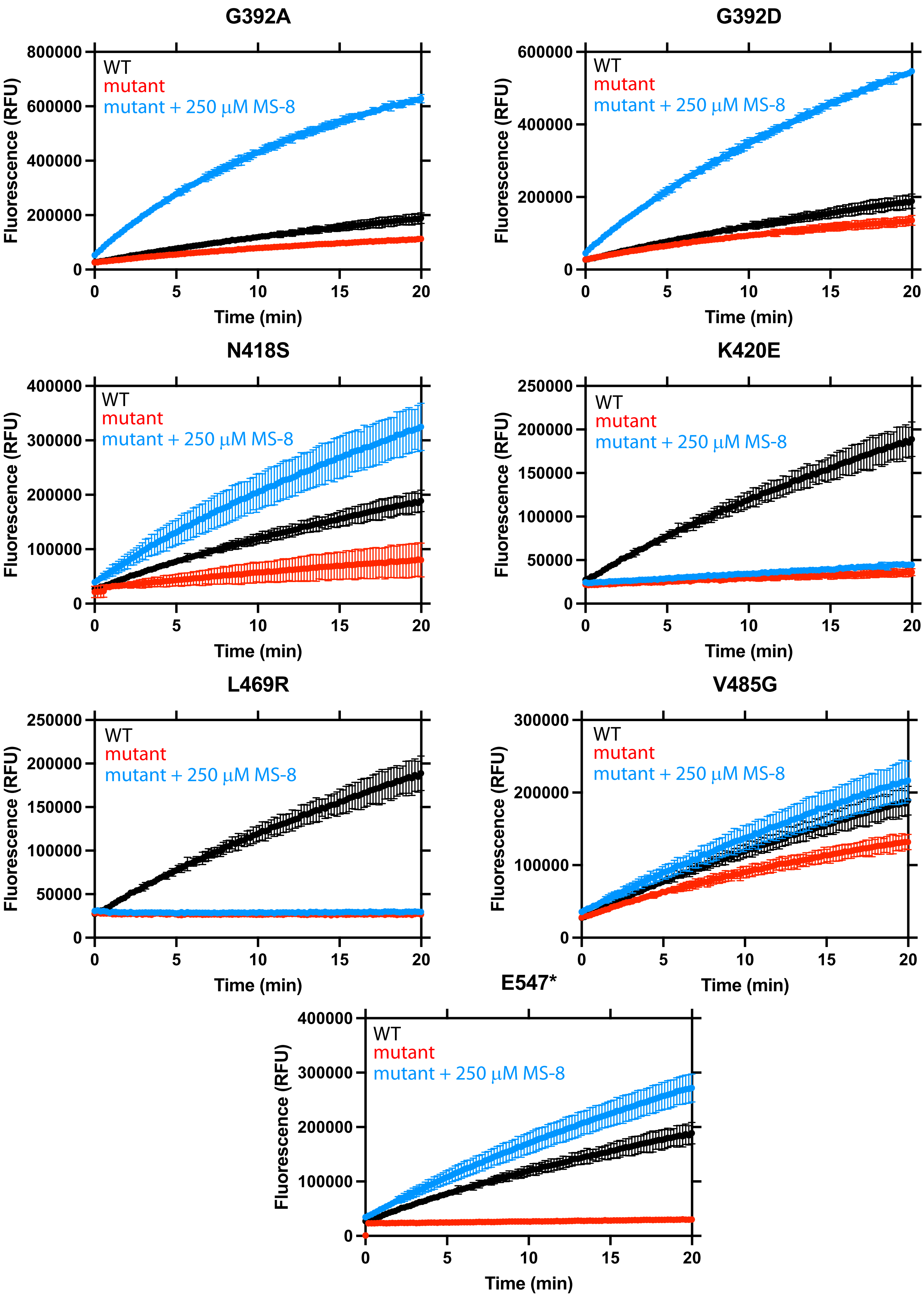

**Figure S4. MS-8 enhances the activity of USP7 variants.**
Comparison of the time course of deubiquitination reaction for FL-USP7 variants alone (red) and treated with 250 μM MS-8 (blue). The untreated WT curve is shown for reference (black). 0.1 nM USP7 was used with 500 nM ubiquitin-rhodamine as its fluorogenic substrate.

Table S1
Summary of enzyme kinetics and ubiquitin-binding affinities of USP7 mutations associated with Hao-Fountain syndrome.

| | WT | M225I | P273H | Y331H | E345K | L373F | G392A | G392D | N418S | K420E | V485G | L469R |
|---|---|---|---|---|---|---|---|---|---|---|---|---|
| **Catalytic domain** | | | | | | | | | | | | |
| $k_{cat}$ ($min^{-1}$) | 1.6±0.0 | 2.5±0.2 | 1.3± 0.2 | 2.4±0.1 | 0.1±0.0 | 0.8±0.0 | 3.6±0.2 | 3.5±0.1 | 10.6±0.7 | N/A | 0.2±0.0 | 0.5±0.0 |
| $K_M$ (μM) | 0.6± 0.1 | 0.6±0.1 | 1.9±0.4 | 0.1±0.1 | N/A | 1.1± 0.2 | 0.2±0.1 | 0.1±0.0 | 0.3±0.1 | N/A | N/A | 0.2± 0.1 |
| $k_{cat}/K_M$ ($min^{-1}\mu M^{-1}$) | 2.9±0.3 | 4.6±1.1 | 0.7±0.2 | 18.6±7.0 | N/A | 0.7±0.1 | 17.8±4.8 | 35.9±11.4 | 30.9±9.2 | N/A | N/A | 2.9±1.1 |
| $K_D$ (μM) | 160.1±24.2 | 91.6±5.2 | 82.5±9.9 | 26.0±7.9 | 621.0±56.6 | 107.9±29.8 | 77.2±4.7 | 33.6±6.6 | 73.4±6.4 | 114.5±8.8 | 171.0±22.6 | N/A |
| **Full-length** | | | | | | | | | | | | |
| $k_{cat}$ ($min^{-1}$) | 56.2±3.4 | 25.5± 2.7 | 19.7±0.7 | 45.3±3.4 | 0.2± 0.0 | 16.1±3.1 | 31.9±1.1 | 30.9±1.5 | 16.1±0.7 | 1.0±0.1 | 1.3±0.1 | 1.8±0.2 |
| $K_M$ (μM) | 0.7±0.1 | 0.5±0.1 | 0.2±0.0 | 0.6±0.1 | N/A | 2.8±0.8 | 0.3±0.0 | 0.3±0.0 | 0.2±0.0 | N/A | N/A | 0.21±0.1 |
| $k_{cat}/K_M$ ($min^{-1}\mu M^{-1}$) | 86.3±14.4 | 55.1±16.4 | 132.0±20.8 | 76.0±15.9 | N/A | 5.9±2.0 | 104±13.1 | 123.1±21.6 | 102.5±20.2 | N/A | N/A | 8.2±3.0 |
